# Targeting KRAS-mutant stomach/colorectal tumours by disrupting the ERK2-p53 complex

**DOI:** 10.1101/2020.08.12.247460

**Authors:** Xiang Wang, Qing Xie, Yan Ji, Jiayan Shen, Yongfeng Zhang, Feng Jiang, Xiangyin Kong, Dandan Liu, Leizhen Zheng, Chen Qing, Jing-Yu Lang

## Abstract

KRAS is widely mutated in human cancers, resulting in nearly unchecked tumour proliferation and metastasis. No therapies have been developed for targeting KRAS-mutant tumours. Herein, we observed that mutant KRAS specifically promoted the formation of ERK2-p53 complex in stomach/colorectal tumour cells. Disruption of this complex by applying MEK1/2 and ERK2 inhibitors elicits strong apoptotic responses in a p53-dependent manner, validated by genome-wide knockout screening. Mechanistically, p53 physically associates with phosphorylated ERK2 through the hydrophobic interaction in the presence of mutant KRAS, which suppresses p53 activation by preventing the recruitment of p300/CBP; trametinib disrupts the ERK2-p53 complex by reducing ERK2 phosphorylation, allowing the acetylation of p53 protein by recruiting p300/CBP; acetylated p53 activates PUMA transcription and thereby kills KRAS-mutant tumours. Our study unveils an important role of the ERK2-p53 complex and provides a potential therapeutic strategy for treating KRAS-mutant cancer via ERK2 inhibition.

## INTRODUCTION

RAS proteins, including HRAS, NRAS, KRAS4A and KRAS4B, act as central regulators of numerous receptor tyrosine kinase (RTK) signalling pathways (Kolch, 2005; Malumbres & Barbacid, 2003; Simanshu et al, 2017). In response to extracellular stimuli, RAS proteins are switched from an inactive GDP-bound state to an active GTP-bound state by recruiting guanine nucleotide exchange factors (GEFs), thereby activating the downstream RAF/MEK/ERK, PI3K-mTOR and RALGDS cascades. The members of the RTK/RAS/RAF/MEK pathway are frequently altered in human cancers, enabling tumours to grow unchecked and metastasize (Downward, 2003). For example, KRAS, together with NRAS and HRAS, is mutated in pancreatic (63.6%), colorectal (37.7%), lung (31.8%), uterine (20%) and stomach (16.4%) adenocarcinomas (Hoadley et al, 2018; Liu et al, 2018b). BRAF is deregulated in approximately 50-60% of melanoma and thyroid adenocarcinomas, while *MAP2K1*and *MAP2K2*, which encode MEK1/2 proteins, are mutated in approximately 4-6% of melanomas. Although RAS proteins play a causal and dominant role in human cancers, it is widely accepted that they are undruggable due to the lack of an accessible site for chemical inhibitors(McCormick, 2018). Newly developed KRAS^G12C^-specific covalent inhibitors exhibit favourable antitumour activity in preclinical settings, accounting for 18.4% of total KRAS mutations at glycine 12(Hoadley et al, 2018), but have no effects on other KRAS^G12^ mutants (Janes et al, 2018; Ostrem et al, 2013). BRAF and MEK1/2 kinase inhibitors, which targets the downstream effectors of RAS, have been approved for treating BRAF-mutant melanoma, achieving remarkable clinical responses (Ascierto et al, 2013; Chapman et al, 2011; Flaherty et al, 2010). But they only have moderate inhibitory effects in KRAS-mutant lung and pancreatic cancers due to cytostatic responses (Jing et al, 2012; Manchado et al, 2016; Solit et al, 2006). In general, the development of therapies that directly target RAS for patient use largely remains an unmet goal (Downward, 2003; McCormick, 2015; McCormick, 2018). It is of great value to develop a strategy for inducing the apoptosis of RAS-mutant tumour cells.

In this study, we reveal that mutant KRAS promotes the formation of ERK2-p53 complex by selectively activating ERK2 phosphorylation in stomach/colorectal tumour cells. The three hydrophobic amino acids of the transactivation domain (TAD) of p53 protein is critical for binding with phosphorylated ERK2. Disruption of ERK2-p53 complex potently induces the apoptosis of KRAS-mutant stomach/colorectal tumour cells, which is achievable by application of MEK1/2 or ERK2 inhibitors. For example, Trametinib, a reversible MEK1/2 kinase inhibitor (Gilmartin et al, 2011), strongly disrupts the ERK2-p53 protein complex by inhibiting ERK2 phosphorylation at threonine 185/tyrosine 187 and promotes the acetylation of p53 protein at lysine 381 and 382 sites by recruiting p300/CBP, thereby activating p53-mediated *PUMA* transcription. Using genome-wide knockout screening and functional validation, we demonstrate that ERK2 inhibition is sufficient to promote the apoptosis of KRAS-mutant tumour cells, allowing the recruitment of p300/CBP to p53, and thereby activates p53-mediated *PUMA* transcription via acetylation. Collectively, our study reveals an important role of the ERK2-p53 complex in maintaining the survival of KRAS-mutant stomach/colorectal tumour cells and provides a novel therapeutic strategy for eliminating these tumours by disrupting the ERK2-p53 interaction.

## RESULTS

### p53 protein is predominantly sequestered by ERK2 in KRAS-mutant stomach/colorectal tumour cells, which is potently disrupted by MEK1/2 inhibitor

Using a coimmunoprecipitation assay, we observed that p53 selectively bound ERK2 in KRAS^G12D^ mutant AGS cells, while ERK1 and other RAS signalling members were not profiled (Figure 1A). Interestingly, the ERK2-p53 complex was strikingly disrupted by MEK1/2 inhibitor trametinib. We obtained similar results in KRAS^G12D^ mutant GSU cells (Figure 1B). Due to the absence of an ERK2-specific immunofluorescent antibody, we used an antibody that recognizes ERK1/2 proteins to detect whether p53 and ERK2 colocalized. Confocal image of double immunofluorescent staining data revealed that the majority of p53 protein was colocalized with ERK1/2 in the cytosol of AGS cells, while trametinib significantly abolished this colocalization (Figure 1C). Reciprocal immunoprecipitation data confirmed that p53 bound ERK2 (Figure 1D). These results suggest that p53 selectively interacts with ERK2 in KRAS-mutant tumour cells, which is disrupted by MEK1/2 inhibition.

**Figure 1.**
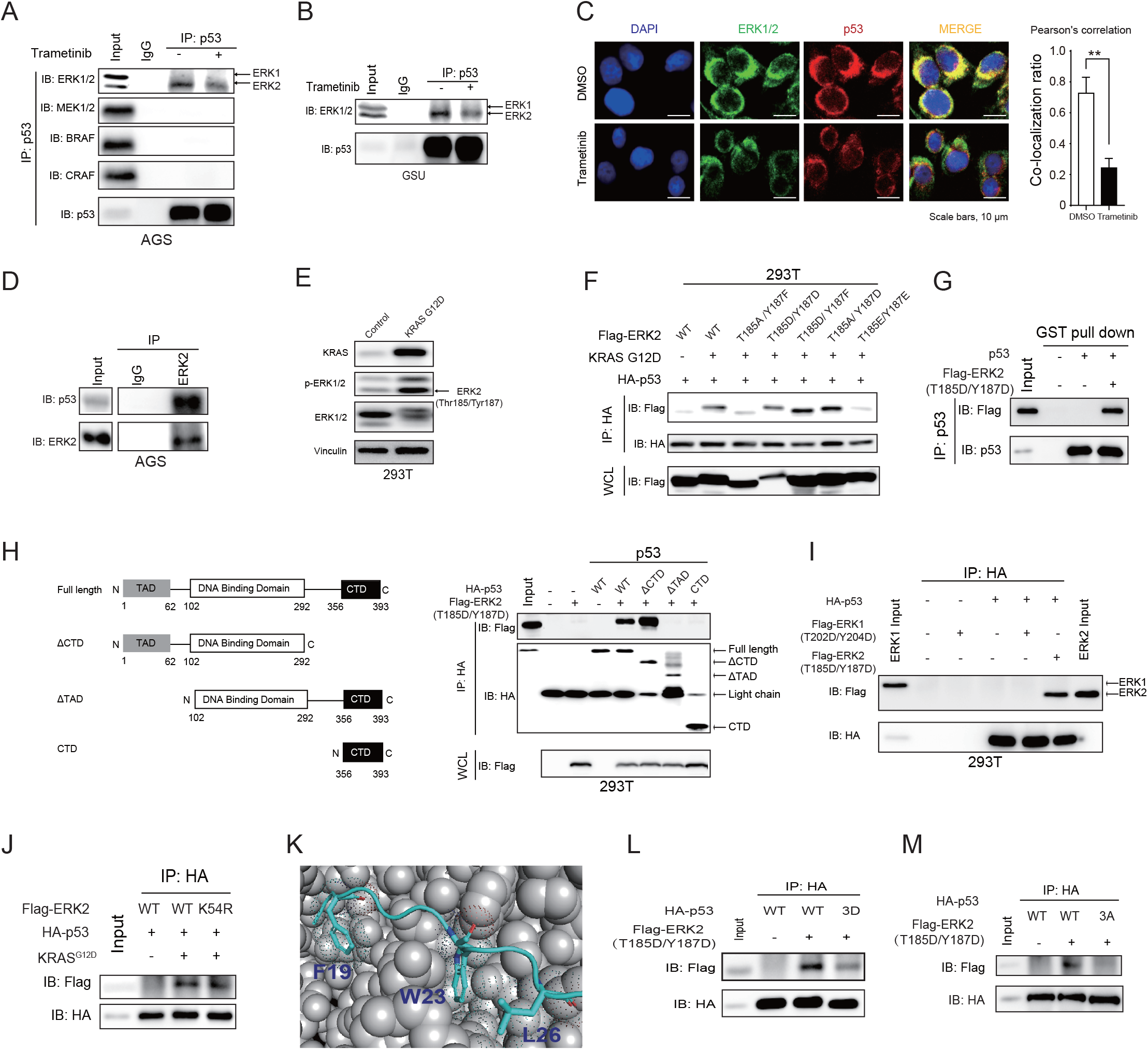
Mutant KRAS promotes the physical interaction between ERK2 and p53 proteins by regulating ERK2 phosphorylation at Thr185 and Tyr187 sites. (A) Coimmunoprecipitation data showed that p53 physically interacted with ERK2 but not ERK1, MEK1/2, BRAF or CRAF proteins in KRAS-mutant AGS cells. The ERK2-p53 complex was largely disrupted by treatment with 1 μM trametinib for 12 hours. (B) p53 physically associated with ERK2 but not ERK1 in KRAS-mutant GSU cells, and this interaction was strongly inhibited by trametinib. (C) Immunofluorescent double staining showed that p53 largely colocalized with ERK1/2 in AGS cells, while trametinib abolished this colocalization. The colocolization of ERK1/2 and p53 was analyzed by Image J. (D) Reverse immunoprecipitation data revealed that ERK2 physically interacted with p53 in AGS cells. (E) Exogenous expression of the KRASG12D mutant robustly enhanced ERK2 phosphorylation (Thr185/Tyr187) in 293T cells. Vinculin was used as a loading control. (F) Exogenous expression of KRASG12D mutant enhanced the physical interaction between ERK2 and p53, and both Thr185 and Tyr187 sites were critical for ERK2 to interact with p53. The four indicated ERK2 mutants, which mimicked double-, single and de-phosphorylated ERK2, were transiently transfected into 293T cells and subjected to coimmunoprecipitation assay after cultured for 48 hours. Wild-type p53 in the absence or presence of the KRASG12D mutant was used as a control. (G) GST pull-down assay for detecting the direct association of purified ERK2 T185D/Y187D and p53 proteins with no ATP. (H-I) The transactivating domain (TAD) is required for p53 to interact with phosphorylated ERK2 but not phosphorylated ERK1 when equal amounts of indicated proteins were loaded. (J) ERK2 K54R mutant had similar binding affinity with p53 protein compared to that of wild-type. (K) Protein-peptide docking analysis of potential amino acids of p53 TAD domain that interact with ATP-bound form ERK2 (PDB:4GT3). (L-M) p53 F19D/W23D/L26D (3D) and F19A/W23A/L26A (3A) mutants abolished their binding affinity with phosphorylated ERK2 compared to that of wild-type.

### Mutant KRAS promotes the formation of the ERK2-p53 complex by phosphorylating ERK2 at the T185 and/or Y187 sites

Next, we asked whether *KRAS* gene mutations can initiate the formation of the ERK2-p53 complex. Exogenous expression of the KRAS^G12D^ mutant obviously increased the levels of phosphorylated ERK2 (Thr185/Tyr187) (Figure 1E) (Botta et al, 2012). More importantly, KRAS^G12D^ mutant visibly increased the physical association between p53 and ERK2 proteins (Figure 1F). To further determine whether ERK2 phosphorylation is required for binding with p53 protein, we generated a series of ERK2 mutants harbouring alanine, aspartic acid or glutamic acid substitutions at threonine 185 and/or tyrosine 187(Prowse et al, 2000). Mutation substitution data revealed that the ERK2 T185A/Y187F mutant, which mimics dephosphorylated ERK2, completely lost its binding affinity with p53 protein, while the T185D/Y187F, T185A/Y187D and T185D/Y187D mutants that mimic single- or double-phosphorylated ERK2 maintained comparable or better binding ability with p53 protein than did wild-type ERK2 (Figure 1F). We did not observe a notable protein association between the ERK2 T185E/Y187E mutant and p53 protein compared with that of wild-type ERK2, suggesting that this mutant is not a good phosphomimetic. Moreover, GST pull-down data revealed that purified ERK2 T185D/Y187D mutant directly associated with purified p53 protein with no ATP addition (Figure 1G). These data suggest that KRAS mutation is a key event for assembling the ERK2-p53 complex by upregulating ERK2 phosphorylation at T185/Y187.

### p53 protein interacts with ERK2 through the hydrophobic amino acids of N-terminal TAD

Next, we generated a series of p53 domain truncation mutants to determine which p53 protein domain is required for binding with phosphorylated ERK2 (Figure 1H left). Domain truncation analysis revealed that the p53 protein lacking the TAD completely lost its binding ability with the ERK2 T185D/Y187D mutant that mimicked double-phosphorylated ERK2, while the p53 ∆CTD mutant exhibited a binding affinity to phosphorylated ERK2 comparable with that of wild-type p53 (Figure 1H right). Using Flag-ERK2 T185D/Y187D as a positive control, we observed no visible physical association between ERK1 T202D/Y204D mutant and p53 when equal amounts of ERK1 T202D/Y204D and ERK2 T185D/Y187D mutant proteins were presented (Figure 1I). These results suggest that p53 protein selectively interacts with ERK2 via the TAD domain.

Despite the kinase activity of ERK2 protein, we observed that there was little or no detectable signal of p53 phosphorylation at eight phosphorylation sites located in the TAD (Supplemental Figure 1)(Yeh et al, 2004). To exclude the role of ERK2 kinase activity, we generated ERK2 K54R kinase dead mutant and observed that ERK2 K54R mutant had similar binding affinity comparable with that of wild-type p53 (Figure 1J), suggesting that the ERK2 kinase activity is dispensable for assembling the ERK2-p53 complex. It was further supported by the fact that both ERK2 T185D/Y187D and T185E/Y187E mutants failed to activate ERK substrate p90RSK phosphorylation (Supplemental Figure 2). This observation led us to investigate whether the protein conformation change of phosphorylated ERK2 but not its kinase activity is essential for forming the ERK2-p53 complex.

Using protein-peptide docking analysis (Zhou et al, 2018), we revealed that three hydrophobic amino acids (Phenylalanine 19, Tryptophan 23 and Leucine 26) of p53 TAD domain were potentially critical for p53 to interact with ERK2 (Figure 1K). To verify this hypothesis, we generated p53 F19A/W23A/L26A (3A) and F19D/W23D/L26D (3D) mutants and observed that both p53 3A and 3D mutants greatly reduced their binding affinity with ERK2 T185D/Y187D mutant compared to that of wild-type p53 (Figure 1L and 1M). These data suggest that p53 physically associates with ERK2 through the hydrophobic interaction.

### MEK1/2 inhibitor potently induces the apoptosis of KRAS-mutant stomach/colorectal cancer cells

Due to the lack of therapeutic agents directly targeting mutant KRAS, we asked whether disruption of the ERK2-p53 complex is able to induce the apoptosis of KRAS-mutant tumours. Herein, we collected seven KRAS mutant stomach/colorectal cancer cell lines and observed that trametinib, a potent and reversible MEK1/2 kinase inhibitor (Gilmartin et al, 2011; Jing et al, 2012), potently reduced the viability of 5 of 7 KRAS-mutant tumour lines with IC_50_ values of 10-30 nM (Figure 2A), suggesting that RAS-mutant stomach/colorectal cancer cells are hypersensitive to MEK1/2 inhibition. The KRAS mutation status of seven tested tumour cell lines was validated by Sanger sequencing (Figure 2B). Immunoblotting data illustrated that trametinib treatment provoke PARP cleavage, a typical signature of apoptosis, in the KRAS-mutant cancer cell lines sensitive to trametinib but failed to do so in two cell lines resistant to trametinib (Figure 2C). However, the bioactivity of MEK1/2 was markedly inhibited by trametinib in all of the tested cancer cell lines regardless of drug sensitivity, as indicated by reduced levels of ERK1/2 phosphorylation (Figure 2C)(Gilmartin et al, 2011). Flow cytometry analysis also revealed that trametinib robustly increased the sub-G1 DNA content by approximately 26% in both KRAS^G12D^ mutant AGS and GSU cells after 48 hours of treatment compared that observed in vehicle-treated cells (p<0.001)(Figures 2D-2G). Noteworthy, trametinib reduced AGS and GSU cell sizes by approximately 50% (p<0.0001, Supplemental Figure 3), resulting 2N and 4N DNA content shifting to the right. These data suggest that disruption of ERK2-p53 complex initiates the apoptosis of KRAS-mutant stomach/colorectal tumour cells.

**Figure 2.**
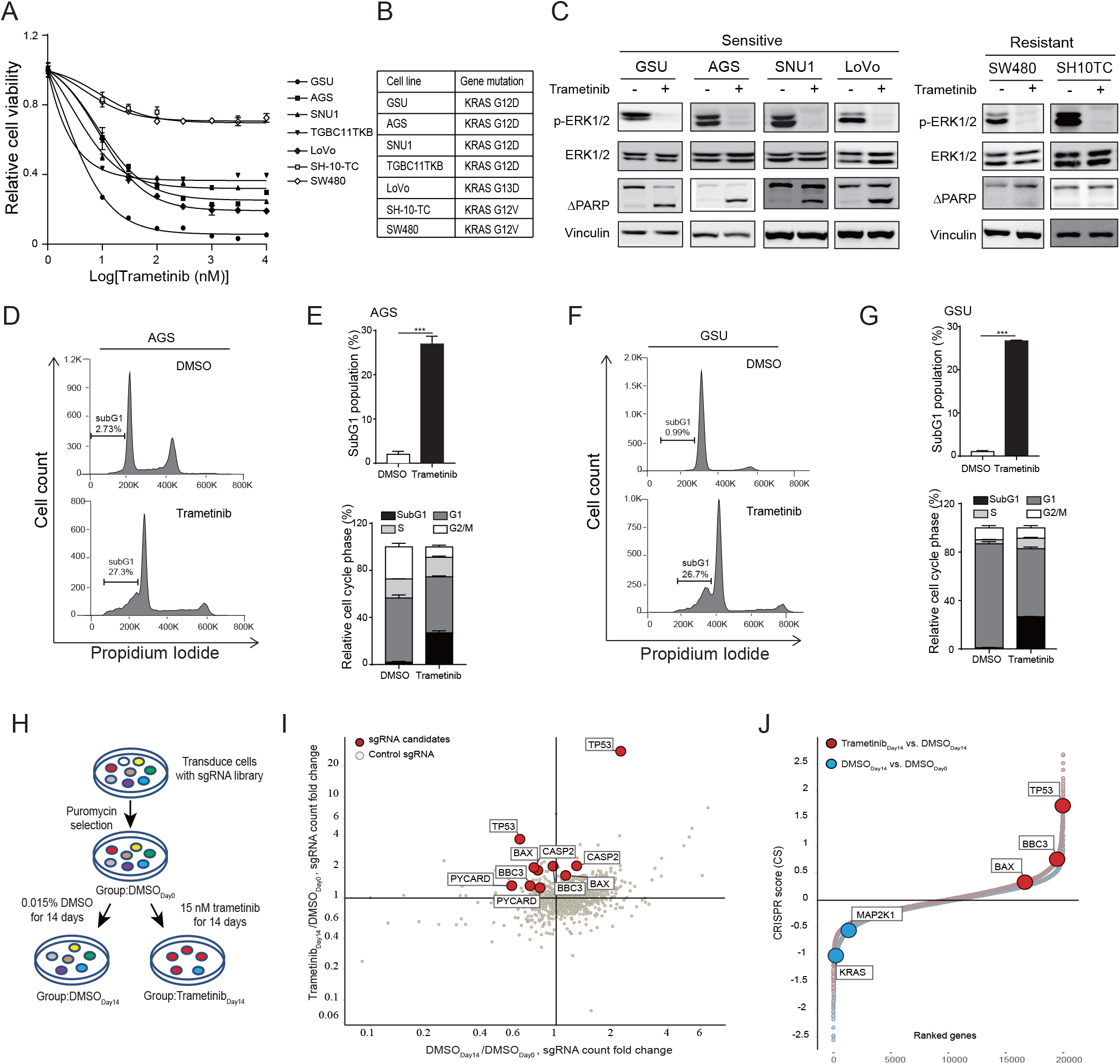
Trametinib potently induces the apoptosis of KRAS-mutant stomach/colorectal cancer cell lines. (A) The viability of 7 KRAS-mutant stomach/colorectal cancer cell lines treated with trametinib for 72 hours at the indicated concentrations were determined by an MTT assay. (B) Sanger sequencing was used to verify the gene mutation status of 7 KRAS-mutant tumour cell lines. (C) The protein expression levels of p-ERK1/2 (ERK1 Thr202/Tyr204, ERK2 Thr185/Tyr187), ERK1/2 and cleaved PARP in four sensitive and two resistant KRAS mutant cell lines were detected after treatment with 100 nM trametinib for 4 or 48 hours. Vinculin was used as a loading control. (D-G) GSU and AGS cells were subjected to flow cytometric analysis after treatment with 100 nM trametinib for 48 hours. The percentages of GSU and AGS cells in different phases of the cell cycle after trametinib treatment were calculated. Data represent the means ± SD of three independent experiments. ***p<0.001, Student’s t-test. (H) The working flow chart of the genome-scale CRISPR-Cas9 knockout screen. (I) MAGeCK analysis revealed that sgRNAs targeting *TP53* and downstream genes such as BBC3/PUMA were significantly reduced in the Trametinib_day14_/DMSO_day0_ vs. DMSO_day14_/DMSO_day0_ group. (J) CRISPR score analysis revealed that *TP53* and BBC3 were positively selected in the Trametinib_day14_ vs DMSO_day14_ group, while KRAS was significantly reduced in the DMSO_day14_ vs DMSO_day0_ group.

### Genome-wide knockout screening validates that *TP53* is a key candidate for initiating trametinib-induced apoptosis of KRAS-mutant stomach/colorectal tumour cells

To comprehensively explore the mechanism by which trametinib induces the apoptosis of KRAS-mutant cells, we performed a genome-scale CRISPR-Cas9 knockout (GeCKO) screen using KRAS^G12D^ mutant GSU cells. As described in the Materials and Methods (Liu et al, 2019), GSU cells were first infected with lentiviruses containing the sgRNA library at a multiplicity of infection (MOI) of 0.4 and then selected for 7 days with puromycin to remove uninfected cells. To achieve 300-fold coverage of sgRNAs, 2-3×10 ^7^ cells were harvested for each group after the treatments were stopped. Infected cells were divided into three groups based on the treatment type: direct genomic DNA extraction (DMSO_day0_ group), 14 days of treatment with 0.015% DMSO (DMSO_day14_ group) or 14 days of treatment with 15 nM trametinib (Trametinib_day14_ group) (Figure 2H). After treatment was discontinued, the genomic DNA of each group was harvested, barcode PCR-amplified and deep-sequenced.

Using the MAGeCK algorithm (Li et al, 2014), we identified that sgRNAs targeting *TP53* and its downstream genes, such as *BAX* and *BBC3/PUMA,* were significantly enriched in the Trametinib_day14_/DMSO_day0_ group compared with the DMSO_day14_/DMSO_day0_ group (Figure 2I). Using CRISPR score analysis (Wang et al, 2015), *TP53*, *BBC3* and *BAX* genes were again positively selected (red curve, Figure 2J). As one of the top selected candidates, the levels of sgRNAs targeting *TP53* gene had increased by 12.6- and 5.9-fold with duplicate hits in the Trametinib_day14_/DMSO_day14_ group (FDR < 2.1E-116, Table 1). These data suggest that the *TP53* gene is critical for trametinib-induced apoptosis of KRAS^G12D^ mutant GSU cells. We obtained similar results in another round of GeCKO screening using MAP2K1^Q56P^ mutant OCUM1 cells(Gannon et al, 2016), in which sgRNAs targeting the *TP53* gene were significantly increased 1.3- and 1.9-fold in the Trametinib_day14_/DMSO_day14_ group (FDR<0.05, Supplemental Figure 4). The low fold change in sgRNA levels was probably attributed to the slow growth rate of OCUM1 cells (doubling time >72 hours).

Noteworthy, our screening data also revealed that the *KRAS* gene is critical for maintaining the survival of KRAS^G12D^ mutant tumour cells, showing that the levels of sgRNAs targeting the *KRAS* gene were significantly reduced in the DMSO_day14_/DMSO_day0_ group (blue curve, Figure 2J). Clonogenicity data further demonstrated that shKRAS knockdown dramatically reduced the numbers of GSU cells compared to that of cells treated with scramble shRNA (p<0.001, Supplemental Figure 5). Based on these results, we hypothesize that the ERK2-p53 complex is critical for maintaining the survival of KRAS-mutant stomach/colorectal tumour cells.

### *TP53* is critical for trametinib-induced apoptosis of KRAS-mutant stomach/colorectal tumour cells *in vitro*

To validate the role of p53, we selected two independent sgRNAs to deplete the *TP53* gene in KRAS^G12D^ mutant GSU and AGS cells. Cell viability data showed that *TP53* gene knockout significantly rescued the viability of GSU cells treated with trametinib compared to that of sgControl cells (p<0.001, Figure 3A). The sgRNA knockout efficiency is shown in the upper panel. Similar results were obtained in another KRAS^G12D^ mutated AGS cell line (p<0.001, Figure 3B). Notably, reintroduction of wild-type p53 by a doxycycline-inducible vector greatly restored the sensitivity of *TP53* knockout GSU and AGS cells to trametinib (p<0.001), which was comparable to that of sgControl cells (Figure 3C), suggesting it’s not an off-target effect. Clonogenicity data also illustrated that *TP53* gene knockout markedly increased the numbers of GSU and AGS cells after trametinib treatment, which was abrogated by re-expressing wild-type p53 (Figures 3D and 3E).

**Figure 3.**
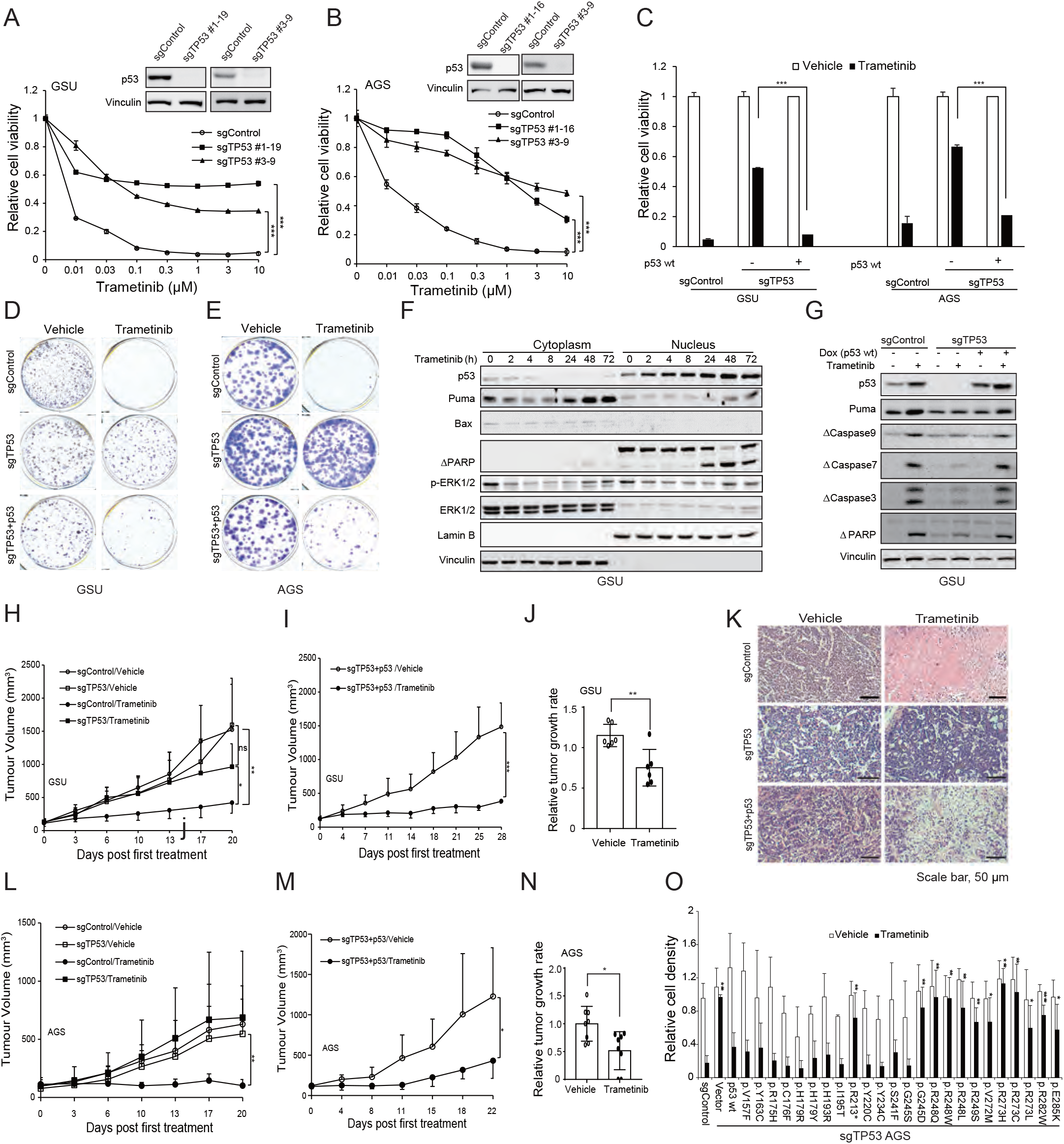
*TP53* is critical for trametinib-induced apoptosis of KRAS-mutant tumour cells. (A) *TP53* gene knockout significantly restored the viability of KRAS-mutant GSU cells. The knockout efficiency of *TP53* gene was shown in the upper panel. (B) *TP53* gene knockout significantly restored the viability of KRAS-mutant AGS cells. The *TP53* gene knockout efficiency was shown in the upper panel. (C) p53 restoration resensitized *TP53*-knockout cells to trametinib in both the GSU and AGS cell lines. (D-E) *TP53* gene knockout significantly increased clonogenic characteristics of GSU and AGS cells, which was abrogated by re-expression of wild-type p53. (F) Trametinib promoted nuclear accumulation of p53 after treatment 100 nM trametinib for 4 hours, peaked at 24 hours, and robustly induced Puma expression and PARP cleavage at 24 hours. Lamin B was used as a nuclear loading control, while vinculin was used as a cytoplasmic loading control. (G) *TP53* knockout significantly blocked trametinib-induced Puma expression, PARP and caspase cleavage in GSU cells, which was reproduced by re-expressing wild-type p53.Vinculin was used as a loading control. (H) Trametinib markedly reduced the in vivo growth of GSU tumour xenografts in nude mice (p<0.01, 3 mg/kg, twice per week), which was overcome by koncking out *TP53* (p>0.05). (I) Ectopic of wild-type p53 resensitized sg*TP53* GSU tumour xenografts to trametinib in athymic mice (p<0.001, 3mg/kg, twice per week). (J) Trametinib induced tumor regression in GSU tumor xenograft model (p<0.01, 3 mg/kg, orally once per day for total three times). (K) Trametinib elicited tumour necrosis of GSU tumour tissue samples as detected by haematoxylin and eosin staining but failed to do so in GSU sgTP53 tumour tissue samples. Tumour necrosis was reproduced by re-expressing wild-type p53. Scale bar, 50 μm. (L) Trametinib markedly reduced the in vivo growth of AGS tumour xenografts in Rag2/Il2r double knckout mice (p<0.01, 3 mg/kg, twice per week), but failed to do so in AGS sgTP53 tumour xenografts (p>0.05). (M) Ectopic of wild-type p53 restored the sensitivity of AGS sgTP53 tumour xenografts to trametinib (p<0.05, 3 mg/kg, twice per week). (N) Trametinib induced tumor regression in AGS tumor xenograft model (p<0.05, 3 mg/kg, orally once per day for total 11 times). The indicated *TP53* mutants were stably transfected into *TP53* knockout AGS cells, and the clonogenicity of these cells was measured after treatment with 1 nM trametinib for 14 days. Data represent the means ± SD of 3 independent experiments. *p<0.05, **p<0.01, ***p<0.001, by Student’s t-test.

Cytoplasmic and nuclear subfractionation data showed that p53 protein accumulated in the nucleus following trametinib treatment for 2 hours and peaked at 24-48 hours, which in turn triggered Puma expression and PARP cleavage in the same time frame (Figure 3F). Bax expression showed little or no changes during the treatment. For better observation of nuclear p53 protein, we loaded equal amounts of cytoplasm and nuclear proteins in these experiments of which the original ratio of cytosol protein versus nuclear protein is approximately 4: 1. To evaluate whether p53 is critical for initiating trametinib-induced apoptosis, we checked the expression of pro-apoptotic proteins in the absence or presence of p53. Immunoblotting data demonstrated that trametinib strongly induced the apoptosis of sgControl GSU cells, as indicated by increased Puma expression and caspase/PARP cleavage (Figure 3G). This phenotype was largely abrogated by knocking out *TP53* but was restored by re-expressing wild-type p53 (Figure 3G). Similar results were obtained in another AGS cell line (Supplemental Figure 6). Thus, *TP53* is a key gene responsible for trametinib-induced apoptosis of KRAS*-*mutant stomach/colorectal tumour cells *in vitro*.

### *TP53* is critical for trametinib-induced tumour growth inhibition of KRAS-mutant stomach/colorectal tumour xenograft models *in vivo*

We further examined whether the *TP53* gene is crucial for transducing the antitumour activity of trametinib *in vivo*. Tumour xenograft data showed that trametinib potently inhibited the *in vivo* growth of GSU tumour xenografts in athymic mice when orally administered at 3 mg/kg twice per week (p<0.01), and this inhibitory effect was largely abolished by *TP53* gene knockout (p>0.05, Figure 3H). Re-expression of wild-type p53 significantly improved the sensitivity of *TP53* knockout GSU tumour xenografts to trametinib compared with vehicle (p<0.001, Figure 3I). Trametinib significantly induced GSU tumour regression at a dose of orally administration at 3 mg/kg once per day (p<0.01, Figure 3J). Immunochemistry staining data showed that trametinib provoked tumour necrosis in sgControl GSU tumour xenografts (Figure 3K), which is consistent with a previous report (Manchado et al, 2016). The tumour necrosis phenotype vanished in the *TP53* knockout group but was restored when wild-type p53 was exogenously expressed (Figure 3K).

Due to the poor growth of AGS tumour xenografts in nude mice, we transplanted the xenografts into Rag2/Il2rg double knockout mice that lack T, B and NK cells. Again, the tumour xenograft data reinforced the notion that trametinib significantly reduced the *in vivo* growth of AGS tumour xenografts (p<0.01), but this antitumour activity was largely abolished by *TP53* gene knockout (p>0.05, Figure 3L). Re-expression of wild-type p53 significantly increased the sensitivity of *TP53* knockout AGS tumour xenografts to trametinib (p<0.05, Figure 3M). Trametinib significantly induced AGS tumour regression at a dose of orally treatment at 3 mg/kg once per day (p<0.05, Figure 3N).

Collectively, these results lead us to conclude that *TP53* is a key gene that determines the therapeutic responses of KRAS-mutant stomach/colorectal tumour xenografts to trametinib *in vitro* and *in vivo*.

### The *TP53* mutation occurs in the DNA binding domain (amino acids 245-285) and the null mutation confers intrinsic resistance of KRAS-mutant stomach/colorectal tumour cells to trametinib

*TP53* is the most frequently deregulated gene in human cancers and overlaps with *KRAS* gene mutations in Pan-cancer Atlas tumour samples (co-occurrence, p<0.001, Supplemental Figure 7A)(Hoadley et al, 2018; Liu et al, 2018b). Herein, we asked which kinds of *TP53* gene mutations affect the therapeutic responses of KRAS-mutant tumour lines to trametinib. We selected the top 24 most frequent *TP53* mutant forms based on the Cosmic and TCGA databases (Cerami et al, 2012; Forbes et al, 2017; Gao et al, 2013) and cloned them into a gene expression vector. The clonogenicity data revealed that point mutations in the p53 DNA binding domain (amino acids 245-285) and nonsense mutations such as TP53^R213∗^ efficiently abolished the therapeutic effects of trametinib *in vitro*, while other hotspot mutations such as *TP53^R175H^* failed to do so (Figure 3O and Supplemental Figure 8). These data explain why SW480 (homozygous TP53^R273C^) and SH-10-TC (heterozygous TP53^R273H/P309S^) cancer cell lines are primarily resistant to trametinib despite the fact they harbour the KRAS^G12V^ mutation (Figure 2A and 2C)(Barretina et al, 2012; Rochette et al, 2005). Collectively, p53 protein, especially the activity of its DNA binding domain, is critical for initiating trametinib-induced apoptosis of KRAS-mutant stomach/colorectal tumour cells.

### ERK2 inhibition promotes the apoptosis of KRAS-mutant stomach/colorectal tumour cells by activating p53-mediated *PUMA* transcription

To determine whether ERK2 inhibition alone is sufficient to eliminate KRAS-mutant tumour cells, we used shRNA to silence ERK2 as well as ERK1 in both *TP53* wild-type and knockout GSU cells. Clonogenicity data showed that shERK2 knockdown significantly reduced the numbers of sgControl GSU cells, a response which mimicked that of trametinib treatment (p<0.001, Figure 4A). *TP53* knockout significantly abrogated the inhibitory effects of both shERK2 knockdown and trametinib (p<0.01, Figure 4A). shERK1 knockdown had little or no effect in both cases, suggesting that this is an ERK2-specific event. Next, we repeated this experiment with two ERK2 kinase inhibitors, AZD0364 and SCH772984 (Pegram et al, 2019; Ward et al, 2019). Cell viability data showed that both AZD0364 and SCH772984 significantly reduced the viability of KRAS^G12D^ mutant GSU cells, but their inhibitory effects were significantly reduced by *TP53* gene knockout (Figure 4B). Similar to trametinib treatment, both AZD0364 and SCH772984 stabilized p53 protein levels, increased Puma expression and promoted PARP cleavage (Figure 4C). p21 and MDM2 expression was also decreased after treatment with trametinib or ERK2 inhibitors (Figure 4C and Supplemental Figure 9). Phosphorylation of the ERK effector p90RSK, which is akin to the inhibitory activity of ERK2 inhibitors (Ward et al, 2019), declined after AZD0364 and SCH772984 treatment. Taken together, our data suggest that ERK2 inhibition is sufficient to promote the apoptosis of KRAS-mutant tumour cells.

**Figure 4.**
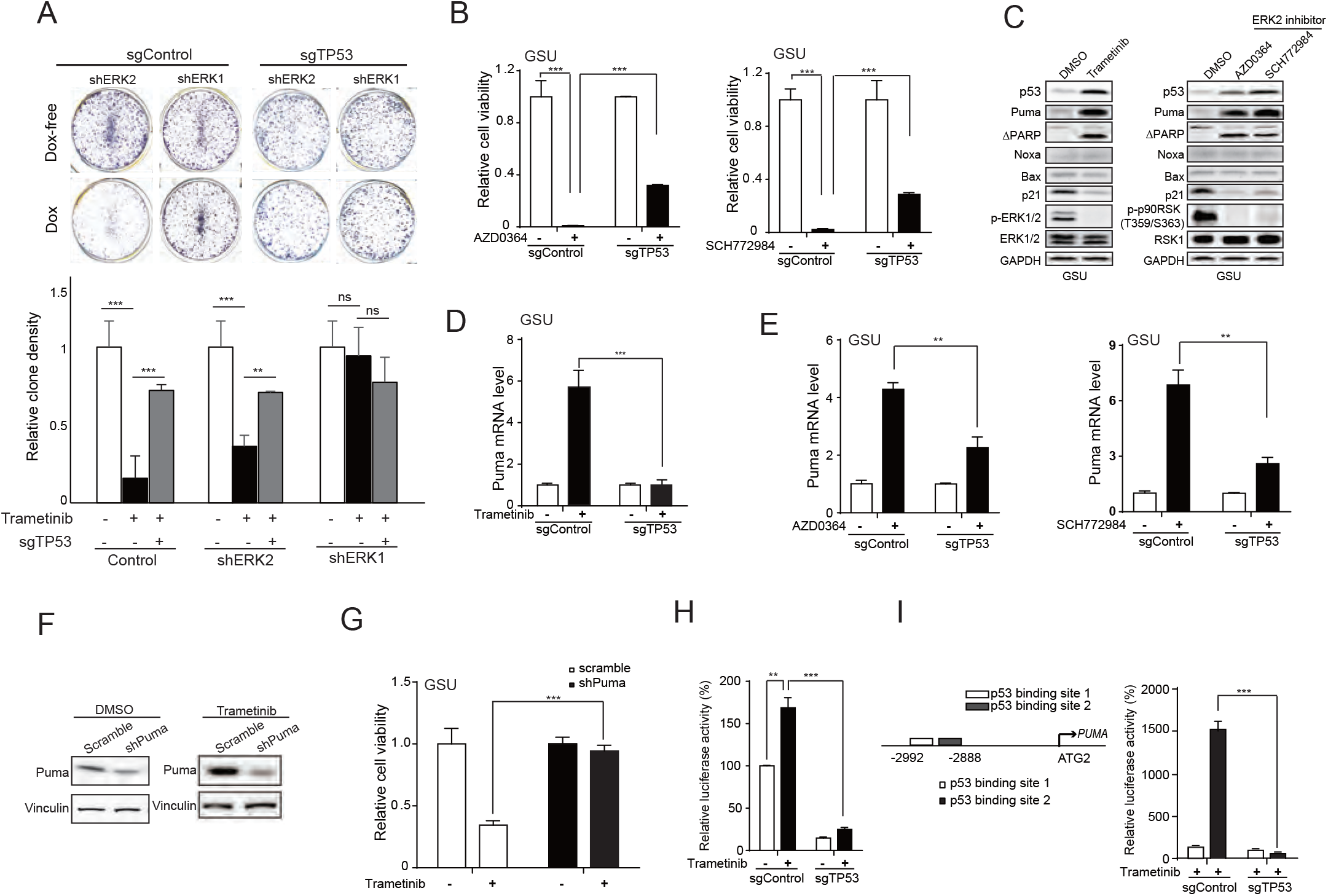
ERK2 inhibition is critical for trametinib-induced apoptosis of KRAS-mutant tumour cells via acetylation of 53 at lysine 382. (A) Knockdown of ERK2 but not ERK1 significantly decreased the colony-forming ability of GSU cells, but this inhibitory effect was abolished by *TP53* gene knockout. 1 nM Trametinib was used as a control. (B)The ERK2 inhibitor AZD0364 and SCH772984 at 1-3 μM potently reduced the viability of GSU sgControl cells but had much less pronounced inhibitory effect in sgTP53 cells. (C) AZD0364 and SCH772984, the effects of which mimic those of trametinib, robustly increased the protein expression of Puma and the cleavage of PARP after treatment for 48 hours but exhibited little or no effect on the expression of Noxa and BAX. The inhibitory effects of AZD0364 and SCH772984 were determined by monitoring the levels of the ERK2 effector phosphorylated p90RSK (Thr359/Ser363). GAPDH was used as a loading control. (D) Trametinib induced Puma mRNA expression in a p53-dependent manner. (E) The ERK2 inhibitor AZD0364 and SCH772984 also elicited Puma mRNA expression in a p53-dependent manner. (F) The shPuma knockdown efficiency in the absence and presence of trametinib was shown in the upper panel. (G) shPuma robustly abolished the inhibitory effect of trametinib in GSU cells. (H) Trametinib induced the luciferase activity of 250-bp PUMA promoter in a p53-dependent manner, and p53 binding site 2 of the PUMA promoter was critical for trametinib-induced PUMA transcription (I).

Among the downstream effectors of p53 signalling, Puma, Noxa and Bax are critical for executing apoptosis (Jeffers et al, 2003; Levine & Oren, 2009; Schuler & Green, 2001; Shibue et al, 2003; Villunger et al, 2003). Immunoblotting data showed that trametinib as well as two ERK2 inhibitors significantly induced Puma protein expression but had little or no impact on Noxa or Bax protein expression (Figure 4C). The qRT-PCR data illustrated that both trametinib and the two ERK2 inhibitors induced Puma mRNA expression in a p53-dependent manner (Figure 4D and 4E). RNA interference data showed that shPuma knockdown, showing more powerful inhibition of Puma protein in the presence of trametinib, almost completely restored the viability of GSU cells treated with trametinib when compared with that of scramble cells (p<0.001, Figure 4F and 4G), suggesting that p53-mediated *PUMA* transcription is critical for trametinib-induced apoptosis of KRAS-mutant tumour cells.

To determine how p53 regulates *PUMA* transcription, we individually cloned two p53 binding sites of the *PUMA* promoter and 250-bp *PUMA* promoter containing two p53 binding sites into a pGL3-basic-luciferase vector (Yu et al, 2001). Luciferase expression data revealed that trametinib induced 250-bp *PUMA* promoter activity in a p53-dependent manner, of which p53 binding site 2 but not binding site 1 responded to trametinib (Figure 4H and 4I). When *TP53* was depleted, binding site 2 was also inactivated. This suggests that trametinib provokes *PUMA* transcription by recruiting p53 protein to p53 binding site 2 of the *PUMA* promoter, which is critical for inducing the apoptosis of KRAS-mutant tumour cells.

### Phosphorylated ERK2 inactivates p53 by preventing the recruitment of p300/CBP, which is rescued by trametinib treatment

p300/CBP, which shares sharing 96% sequence identity, also interacts with p53 through the F19 and W23 hydrophobic amino acids of N-terminal TAD (Ferreon et al, 2009). Thus, phosphorylated ERK2 potentially blocks the recruitment of p300/CBP to the p53 protein by occupying these hydrophobic amino acids and thereby preventing the activation of the p53 protein via acetylation. Coimmunoprecipitation data revealed that p53 physically associated with ERK2 in the absence of trametinib but preferentially bound p300 in the presence of trametinib (Figures 1A, 1B, 5A and 5B). To support our notion, we performed a competitive assay and observed that exogenously overexpressed ERK2 T185D/Y187D mutant robustly reduced the physical association between p53 and p300 (Figure 5C). Noteworthy, ERK2 T185D/Y187D mutant was no longer able to disrupt the interaction between p300/CBP and p53 when three key hydrophobic amino acids of wild-type p53 were substituted by aspartic acids (Figure 5D), reinforcing the fact that these three hydrophobic amino acids of p53 N-terminal TAD are critical for assembling the ERK2-p53 complex.

**Figure 5.**
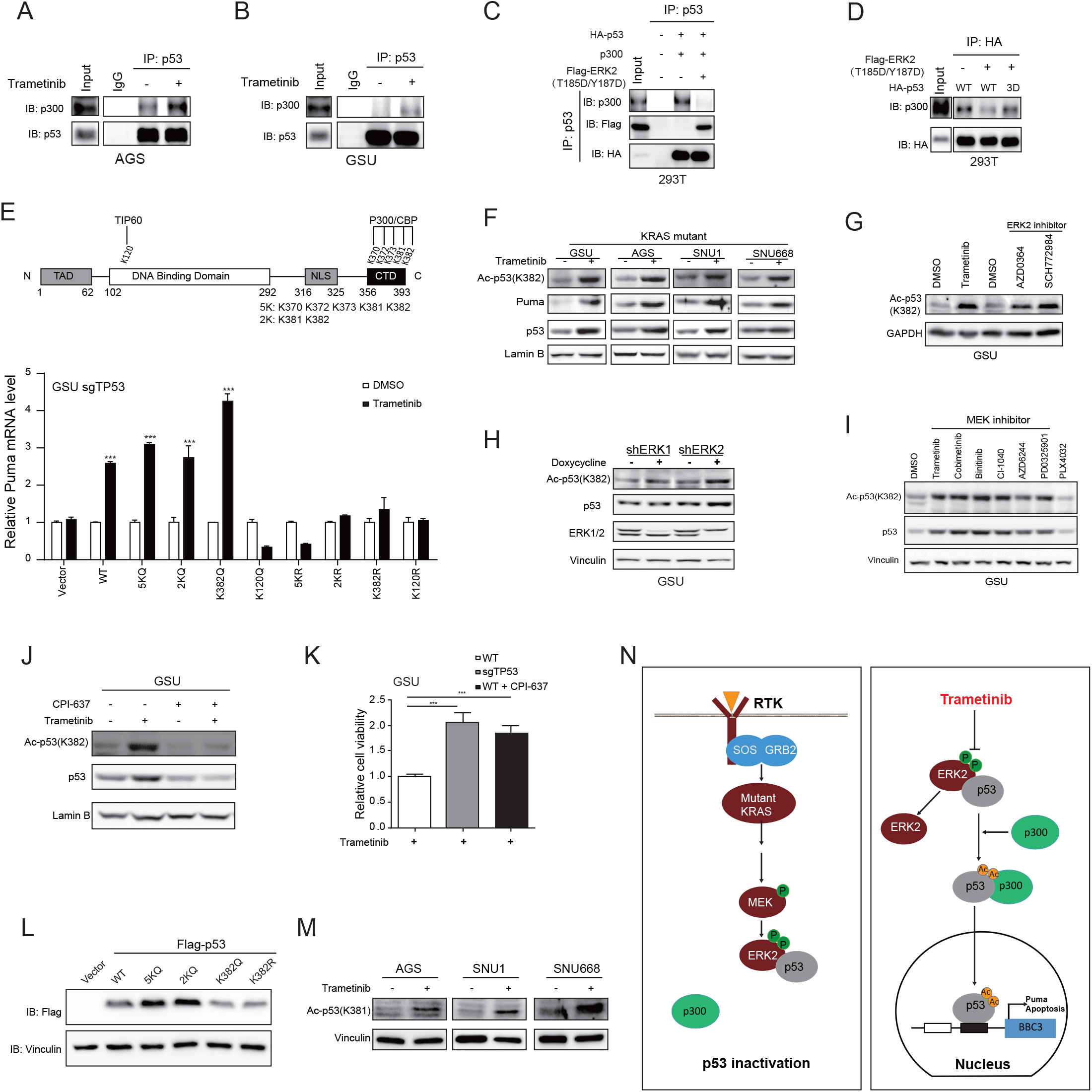
Trametinib promotes p53 acetylation by recruiting p300/CBP. (A-B) Trametinib robustly increased the physical assocation between p53 and p300 in KRAS-mutant AGS and GSU cells after treatment for 4-12 hours. (C) Indicated proteins were detected by immunoblotting after immunoprecipitated with anti-p53 antibody when p53 and p300 was co-expressed in the absence and presence of ERK2 T185D/Y187D mutant. (D) Indicated protein were immunoblotted after precipitated with HA-p53 when ERK2 T185D/Y187D mutant was co-expressed with p53 or p53 3D mutant. (E) The p53 wild-type, 5KQ, 2KQ and K382Q mutants, but not other mutants, significantly promoted the mRNA expression of Puma upon trametinib treatment. 5K indicates the lysines at 370, 372, 373, 381 and 382, while 2K indicates the lysines at 381 and 382. The domain structure of the p53 protein was shown in the upper panel. (F) Trametinib increased the levels of K382 acetylation of p53 protein as well as the expression of Puma protein in multiple KRAS-mutant stomach cancer cell lines. Lamin B was used as a loading control. (G) The ERK2 inhibitor AZD0364 and SCH772984 increased the expresssion of K382-acetylated p53 protein. (H) shERK2 but not shERK1 increased the protein expression of total and K382-acetylated p53 in GSU cells. (I) Trametinib with another five MEK1/2 kinase inhibitors with different chemical structures increased both total p53 and K382-acetylated p53 protein expression, while PLX4032 exerted little or no changes. (J) The p300/CPB inhibitor CPI-637 significantly reduced the levels of K382-acetylated and total p53 protein. (K) CPI-637 robustly rescued the viability of KRAS-mutant GSU cells upon trametinib treatment. (L) p53 5KQ and 2KQ mutants but not the K382Q mutant had better protein stability than wild-type. (M) Trametinib increased the acetylation of p53 protein at lysine 381 in multiple KRAS-mutant cell lines. (N) A cartoon of our working model.

In addition, we observed little or no ubiquitination change on p53 protein (Supplemental Figure 10), which is probably caused by weakly or reduced MDM2 expression after trametinib treatment (Supplemental Figure 9). Collectively, our data suggest that the ERK2-p53 complex is predominant in KRAS-mutant stomach/colorectal tumour cells, which prevents the recruitment of p300/CBP to p53 protein.

### The K382 acetylation of p53 protein is crucial for trametinib-induced *PUMA* transcription

In the literature, p300/CBP can simultaneously acetylate p53 at five lysine sites (370, 372, 373, 381 and 382) (Gu & Roeder, 1997; Sykes et al, 2006). To evaluate which acetylation site is critical for p53 transcriptional activity, we constructed a series of p53 mutants harbouring glutamine or arginine substitutions at lysine 120, 370, 372, 373, 381 and 382; these substitutions mimic acetylated or deacetylated lysine, respectively (Figure 5E, upper panel). p53 K120Q mutant acetylated by TIP60 was used as a negative control (Tang et al, 2006). After exogenous expression of the p53 mutants in *TP53* knockout GSU cells, the 5KQ, 2KQ and K382Q mutants, but not K120Q or the other mutants, significantly enhanced Puma mRNA expression levels upon trametinib treatment; the levels were the same as or better than those induced by wild-type p53 (p<0.01, Figure 5E). Of note, the K382Q mutant itself, which mimics the K382-acetylated form of p53 protein, was sufficient to initiate trametinib-induced *PUMA* transcription. Immunoblotting data confirmed that p53 protein was acetylated at lysine 382 after trametinib treatment in multiple sensitive cancer cell lines, including GSU and AGS, accompanied by Puma expression (Figure 5F). Similar to trametinib, AZD0364 and SCH772984 obviously increased p53 and K382-acetylated p53 protein expression levels (Figures 4C and 5G). As expected, shERK2 but not shERK1 increased the protein expression levels of p53 and K382-acetylated p53 (Figure 5H). To examine whether this increase is a general mechanism for MEK1/2 inhibitors, we selected another five MEK1/2 inhibitors with different chemical structures and observed that they all enhanced the protein expression of total p53 and K382-acetylated p53 (Figure 5I). As a negative control, BRAF inhibitor PLX4032 failed to do so.

To further examine whether p300/CBP was required for acetylating p53 protein, we chose a p300/CBP-specific inhibitor CPI-637 and observed that CPI-637 treatment greatly decreased the levels of total and acetylated p53 upon trametinib treatment (Figure 5J). Importantly, CPI-637 treatment robustly increased the viability of GSU cells treated with trametinib to levels comparable to *TP53* knockout (Figure 5K). These results suggest that p300/CBP is critical for acetylating p53 protein at lysine 382, which is a key event for inducing trametinib-mediated apoptosis of KRAS-mutant tumour cells.

It is noteworthy that K382Q and K382R mutants had little or no changes in the p53 protein stability, while 5KQ and 2KQ mutants strongly enhanced p53 protein expression compared with that of wild-type p53 (Figure 5L). The levels of p53 K381 acetylation were visibly increased in trametinib-treated cases (Figure 5M), suggesting that the acetylation of p53 protein at lysine 381 is important for stabilizing p53 protein.

In summary, p53 protein is sequestered by phosphorylated ERK2 through the hydrophobic interaction in KRAS-mutant stomach/colorectal tumour cells, which prevents its association with p300/CBP; trametinib disrupts the ERK2-p53 complex via inhibiting ERK2 phosphorylation, allowing the acetylation of p53 protein at lysine 381 and 382 by recruiting p300/CBP, and eventually promotes p53-mediated *PUMA* transcription and the apoptosis cascade (Figure 5N). Thus, we provide a potential therapeutic strategy for eliminating RAS-dependent cancer by disrupting the ERK2-p53 complex, which is helpful for developing cancer therapies to target diseases with mutant KRAS.

## DISCUSSION

In this study, we revealed the existence of the ERK2-p53 protein complex in KRAS-mutant stomach/colorectal tumour cells, which is specifically induced by mutant KRAS. The physical association of ERK2-p53 complex is highly hydrophobic-dependent, of which the three hydrophobic amino acids (F19, W23 and L26) of p53 TAD domain are essential for forming ERK2-p53 complex. Interestingly, these hydrophobic amino acids are also critical for assembling the p300/CBP-p53 and MDM2-p53 complexes (Feng et al, 2009; Kussie et al, 1996; Teufel et al, 2007). In our case, we showed that phosphorylated ERK2 had much higher binding affinity with wild-type p53 protein compared to p300/CBP, but failed to disrupt the association between p300/CBP and p53 when these three hydrophobic amino acids were substituted by aspartic acids (Figure 5D). The association of MDM2 and p53 is also very weak in KRAS-mutant stomach/colorectal tumour cells, showing no ubiquitination of p53 (Supplemental Figure 10). Thus, we believe that the ERK2-p53 complex is predominant in KRAS-mutant stomach and colorectal tumours, which explains why disruption of this complex is lethal for KRAS-mutant tumour cells.

Noteworthy, acetylated p53 protein is highly selective for regulating Puma transcription. In our case, trametinib selectively induced Puma expression, accompanied with reduced p21 and MDM2 expression (Figure 4C and Supplemental Figure 9). In contrast, DNA damage agents such as mitomycin C often activates p21 expression by phosphorylating p53 at serine 15, but has little or no induction on Puma and MDM2 (Supplemental Figure 11). Unlike wild-type p53 protein, p53 3D, 3A and 2KQ mutants were not responding to mitomycin C after re-introduced back into sg*TP53* GSU tumour cells (Supplemental Figure 11). These data suggest that acetylated p53 protein has a distinct role for regulating Puma expression, while phosphorylated p53 is critical for p21 expression. It also explains why trametinib reduced p21 expression due to the majority of p53 protein was switched to acetylated form.

In the literature, oncogenic Ras can induce p53 accumulation via PML and promotes premature senescence in primary normal cells (Ferbeyre et al, 2000; Pearson et al, 2000; Ries et al, 2000; Serrano et al, 1997). However, trametinib treatment reduced PML expression in KRAS-mutant GSU cells (Supplemental Figure 9C). We detected little or no PML expression in KRAS-mutant AGS cells. Moreover, PML negatively correlated with patient survival at mRNA level (Log-Rank p=0.016, mRNA-high, n=182, mRNA-low, n=172) in TCGA Pan-cancer stomach adenocarcinomas, suggesting that PML is often inactively mutated and lowly expressed in tumour cells (Guan & Kao, 2015). Thus, we believe that PML-mediated p53 accumulation is likely not involved in our case.

The role of ERK2-p53 complex is likely more predominant in stomach/colorectal tumour cells. After analysed the cell viability data of 771 human cancer cell lines obtained from the Genomics of Drug Sensitivity in Cancer database (Yang et al, 2013), we revealed that 127 human cancer cell lines harbouring RAS hotspot mutations at glycine 12, glycine 13 and glutamine 61 were much more sensitive than their 644 wild-type counterparts (p<1.62E-07, Supplemental Figure 12). Remarkably, stomach/colorectal cancer cell lines harbouring RAS mutations were more sensitive to trametinib than RAS-mutant lung, pancreatic and other cancer cell lines. The median IC_50_ values of stomach/colorectal cancer cell lines were 0.3±0.45 μM from a total of 23 tumour lines (p≤0.01), and 9 of them had IC_50_ values less than 36 nM. Recent clinical trials illustrate that trametinib has potential therapeutic value for treating KRAS-mutant colorectal tumours (Huijberts et al, 2020; Wu et al, 2020). Therefore, it would be of great value to examine whether trametinib can treat KRAS-mutant stomach/colorectal tumours in clinical settings.

It is unclear why trametinib exhibited better inhibition in KRAS-mutant stomach/colorectal tumours. Up to now, over 250 substrates downstream of ERK1/2 have been identified (Eblen, 2018; Unal et al, 2017), which corresponds to profound diversity of ERK2-associated complexes. p53-binding proteins are also highly dynamic and are associated with Bcl-xL protein in normal cells (Chipuk et al, 2004; Mihara et al, 2003). Thus, the assembly of the ERK2-p53 complex is likely selective and predominant in KRAS-mutant stomach/colorectal cancers, whose bioactivity is potentially compensated by other ERK2-associated complexes in other cancer subtypes. More efforts are needed to determine the detailed mechanism in the future.

*KRAS* and *TP53* are two of the most frequently mutated genes in human cancer. Thus, *TP53* gene mutations potentially attenuate the therapeutic responses of KRAS-mutant stomach and colorectal adenocarcinomas to MEK1/2 inhibitors (Figure 3O). After analysed with stomach and colorectal cancer samples in public available databases, we observed that *TP53* gene mutations were almost equally distributed in both KRAS-mutant and wild-type stomach and colorectal tumour samples (p>0.05, Supplemental Figure 7B and 7C). *TP53* mutant cases were even less than TP53 wild-type cases in KRAS-mutant metastatic colorectal tumour samples (p<0.0001, Supplemental Figure 7D) (Yaeger et al, 2018). These data indicates that at least half of stomach and colorectal cancer patients harbouring KRAS mutations potentially benefit from MEK1/2 inhibitor treatment. Nevertheless, it is important to check *TP53* mutations to achieve the best therapeutic responses before applying MEK1/2 inhibitors to patients.

Stomach and colorectal adenocarcinomas frequently present amplified expression of RTK/RAS pathway members, such as *MET, FGFR2, EGFR* and *KRAS* (Chia & Tan, 2016; Liu et al, 2018b), all of which can potentially induce the ERK2-p53 complex by activating the MEK1/2-ERK2 cascade. To examine this hypothesis, we collected a total of forty-three stomach cancer cell lines and recorded their MEK1/2 phosphorylation levels under serum-free conditions, as shown in Supplemental Figure 13A (lower right panel, grey curve). After screening, we observed that trametinib efficiently decreased the viability of 18 of 43 stomach cancer cell lines with IC_50_ values of 5-100 nM. Upon retrieving data from the CCLE database (Barretina et al, 2012), we identified that stomach cancer cell lines harbouring KRAS (7/8), NRAS (1/1) and MAP2K1 (1/1) mutations, except SH-10-TC (heterozygous *TP53*^R273H/P309S^), were hypersensitive to trametinib. To our surprise, trametinib exhibited potent inhibitory effects in MET-amplified (4/5) stomach cancer cell lines but not in other RTK-amplified stomach cancer cell lines, with IC_50_ values of 10-50 nM, showing increased levels of Puma and K382-acetylated p53 protein (Supplemental Figure 13A and 13B). Hs746T, which contains a *TP53*^K319*^ mutation, is the only tested MET-amplified cell line with no response to trametinib(Barretina et al, 2012). Re-introduction of wild-type p53 greatly sensitized Hs746T cells to trametinib, which was accompanied by increased Puma expression (p<0.001, Supplemental Figure 13C and 13D). These data suggest that MET amplification is another driving factor for assembling the ERK2-p53 complex.

It is worth mentioning that the BRAF inhibitor vemurafenib/PLX4032, which targets the KRAS downstream effector BRAF(Bollag et al, 2012), had no inhibitory effects on any of the seven tested KRAS-mutant stomach/colorectal cancer cells (Supplemental Figure 14A). To figure out the mechanism, we performed immunoprecipitation assay and observed that PLX4032 did not disrupt the ERK2-p53 complex due to its inability to inhibit ERK2 phosphorylation (Supplemental Figure 15B and 15C). It explains why PLX4032 does not inhibit KRAS-mutant tumour cells (Joseph et al, 2010). However, ERK2 kinase inhibitors strongly inhibited the growth of KRAS-mutant stomach/colorectal cancer cells, which was comparable to the activity of trametinib. Moreover, ERK reactivation is a key event for cells to develop acquired resistance to BRAF and MEK1/2 kinase inhibitors (Ahronian et al, 2015; Komatsu et al, 2015; Little et al, 2013; Liu et al, 2018a). Thus, therapies directly targeting ERK2 may provide a new opportunity for killing KRAS-mutant stomach/colorectal cells with the advantages of circumventing ERK reactivation and reducing the toxicity of ERK1 inhibition.

## METHODS

### Cell Lines

The gastric cancer cell lines were purchased from Korean Cell Line Bank, RIKEN BRC Cell Bank or JCRB Cell Bank, respectively. LoVo and SW480 cell lines were purchased from the cell bank of Shanghai Institutes for Biological Sciences (Shanghai, China). Cells were cultured in either RPMI 1640 or DMEM/F12 medium with 10% fetal bovine serum (Hyclone) and 1% penicillin streptomycin (Life Technologies), and were incubated at 37°C with 5% CO_2_. All cell lines were recently authenticated with STR assays, keeping as mycoplasma-free.

### Reagents

Trametinib, AZD6244 and MG132 were purchased from Selleck Chemicals (Shanghai, China). AZD0364 was purchased from MedChemExpress (MCE) (Shanghai, China). CPI-637 and SCH772984 was purchased from TargetMol. Puromycin, choloroquine, polybrene, NaCl, NaF, Na3VO4, PMSF, Tween 20, sucrose, PEG400, Cremophor® EL, PEG8000 and DMSO were purchased from Sigma-Aldrich (Saint Louis, USA). Thiazolyl blue tetrazolium bromide (MTT) and Tris were purchased from VWR (Radnor, USA). The primary antibodies against phospho-p44/42(ERK1 Thr202/Tyr204, ERK2 Thr185/Tyr187) (#4377, 1:6,000), ERK1/2 (#9102, 1:6,000), acetyl-p53 (Lys382) (#2525,1:2,000), phosphor-p53 (Ser6) (#9285T, 1:1,500), phosphor-p53 (Ser9) (#9288T, 1:1,500), phosphor-p53 (Ser15) (#9286T, 1:1,500), phosphor-p53 (Thr18) (#2529T, 1:1,500), phosphor-p53 (Ser20) (#9287T, 1:1,500), phosphor-p53 (Ser46) (#2521T, 1:1,500), phosphor-p53 (Thr81) (#2676T, 1:1,500), cleaved caspase-3 (#9664, 1:2,000), cleaved caspase-6 (#9761, 1:2,000), cleaved caspase-7 (#8438, 1:2,000), cleaved caspase-9 (#7237, 1:2,000) and cleaved PARP (#5625, 1:1,000) were obtained from Cell Signaling Technology. Primary antibodies against Puma α/β (sc-28226, 1:2,000), Bax (sc-7480, 1:2,000), p53 (sc-126, 1:3,000), p21 (sc-397, 1:3,000), Noxa (sc-56169, 1:1,000), ERK2 (sc-1647, 1:2,000), p300 (sc-48343, 1:3000), lamin B (sc-6216, 1:3,000), HA-probe (sc-7392, 1:2,000) and vinculin (sc-25336, 1:3,000) were obtained from Santa Cruz Biotechnology. Primary antibodies against β-tubulin (ab135209, 1:5,000), phospho-p53 (Thr55) (ab183546, 1:1,500) and acetyl-p53 (K381) (ab61241, 1:3,000) were purchased from Abcam. Primary antibodies against Flag-tag (F3165, 1:3,000) was obtained from Sigma-Aldrich.

### Plasmids

Human CRISPR Knockout Pooled Library (GeCKO v2) (#1000000048), LentiCRISPR v2 (#52961), pMD2.G (#12259), pRSV-Rev (#12253) and pMDLg/pRRE (#12251) were purchased from Addgene. The *TP53* and ERK2 constructs were obtained from CAS_Key Laboratory of Tissue Microenvironment and Tumour. The KRAS^G12D^ and MAP2K1^Q56P^ mutants were cloned from GSU and OCUM1 cells, respectively, validated by Sanger Sequencing. The DNA sequence of 3× p53 binding site 1 (5’-CTCCTTGCCTTGGGCTAGGCC-3’) and 3× p53 binding site 2 (5’-CTGCAAGTCCTGACTTGTCC-3’) was individually cloned into a pGL3-basic luciferase vector. *TP53* and ERK2 mutant constructs were generated by PCR cloning as mentioned. Primers for generating gene were listed in Supplementary Table 2 and shRNA constructs were listed in Supplementary Table 3.

### Cell viability assay

Cells were seeded into 96-well plates at a density of 5,000-6,000/well, and were treated with Chemical at final concentrations of 0.01, 0.03, 0.1, 0.3, 1, 3, and 10 μM on the next day, followed by 72 hours’ incubation at 37°C with 5% CO_2_. When treatment stopped, cells were then added with 20 μl MTT solution for 4 hours, followed by 12-16 hours’ incubation with 50 μl triplex solution (0.012 M HCl, 10% SDS, and 5% isobutanol) before detecting OD570 nm.

### Colony formation assay

Cells were plated into 6-well plates at a density of 1,000-3,000/well, and were treated by trametinib or vehicle (0.3% DMSO) at 37°C for two weeks. After treatment stopped, colonies were fixed and stained with a mixture of 6% glutaraldehyde and 0.5% crystal violet for 1 hour. The density of colonies was captured and analyzed by Image J software.

### Immunoprecipitation

Cells were lysed on ice with 1×#9803 lysis buffer containing 20 mM Tris-HCl (pH 7.5), 150 mM NaCl, 1 mM Na2EDTA, 1 mM EGTA, 1% Triton X100, 2.5 mM sodium pyrophosphate, 1 mM beta glycerophosphate, 1 mM Na_3_VO_4_, 1 mM NaF, 1 mM PMSF and protease inhibitor cocktail (Cat.11873580001, Roche). After pre-cleaned, primary antibodies were added to the supernatant of cell lysates and incubated at 4 °C overnight. The protein A/G agarose beads (SC-2003, Santa Cruz) or Dynabeads^TM^ protein G beads (#10004D, Invitrogen) were added into each sample and incubated at 4 °C for 1-2 hours on the next day. The precipitants were washed with 1× lysis buffer for five times, boiled for 10 min and were subjected to immunoblotting.

### GST pull-down

GST-fused ERK2 T185D/Y187D mutant and p53 wild-type plasmids were transiently transfected into BL21 competent cell. After vigorous shaking (250 rpm) for 12-15 h at 37 °C, the culture medium was diluted with 1:100 ratio and cultured with an A600 of 0.6-0.8 (about 3-5 h) with vigorous agitation at 30 °C. After treated with 0.5 mM IPTG, culture medium were continuously incubated for 4 h at 30 °C. The cell pellets were spin down at 4, 000 rpm for 30 min, washed with1×ice-cold PBS and lysed with 1×#9803 cell lysis buffer with 10-15 times sonication. Each 1 mg protein were incubated with 10 μl pre-washed GST beads with gentle agitation at room temperature for 1 h. After wash with 1× #9803 cell lysis buffer (500 μl 1x 9803 buffer for total 80 μl beads) for three times, GST fusion proteins were eluted by 10 mM or 20 mM reduced glutathione at RT for 30 min. Supernatants were transferred to fresh tubes after centrifuged at 1,000 g for 5 min. For GST pull-down assay, 250 μg purified GST-p53 protein and 250 μg GST-ERK2 T185D/Y187D mutant protein were mixed, incubated with 0.2 μg anti-p53 antibody with gentle agitation at 4 °C overnight. p53-associated protein were pulled down by 5 μl Dynabeads^TM^ protein G beads (#10004D, Invitrogen) at 4 °C for 1-1.5 h. The precipitants were washed with 1× #9803 cell lysis buffer for five times, boiled for 10 min and subjected to immunoblotting.

### Quantitative PCR

Total RNA was extracted from cultured cells using Trizol^TM^ reagent, and was reversed into cDNA with PrimerScript^TM^ RT reagent Kit (RR037A). Quantitative PCR was performed with NovoStart SYBR qPCR supermix using ABI-7500 instrument. GAPDH was used as an internal reference to normalize input cDNA. Primers for real time PCR were listed in Supplementary Table 4.

### CRISPR/Cas9-mediated knock out

Two annealed sgRNA oligos targeting *TP53* gene were cloned into lentiCRISPRv2 vector after digested with BsmBI. Target cells were transiently transfected with sgRNA plasmids using Lipofectamine® 3000, followed by puromycin selection for 3 days to remove uninfected cells. Cells were seeded into 96-well plates at a density of one cell per well, and knockout clones were validated by immunoblotting and Sanger sequencing. sgRNA oligos were listed in Supplementary Table 3.

### Immunofluorescence staining

Cells were plated into 24-well plates pre-seated German glass cover slips, and were subjected to trametinib or vehicle (0.3% DMSO) for indicated time. After fixed with 4% formaldehyde, cells were blocked with 3% BSA for 1 hour and incubated with primary antibody at 4 °C overnight. After wash with three times, cells were incubated with Alexa Fluor® 488 or Rhodamine Red^TM^-X labeled secondary antibody (Jackson ImmuoResearch) at room temperature for 1 hour. DAPI was added into the cells before mounting and cells were imaged with a fluorescence microscope.

### Immunohistochemistry staining

After dissecting tumour tissues, samples were fixed with 4% paraformaldehyde at 4°C overnight. Then, tumour samples were stained with haematoxylin and eosin staining (Wuhan Servicebio Technology, China) following standard protocol.

### Cytoplasmic and Nuclear Extraction

Cells were first lysed with 800 μl lysis buffer (10 mM HEPES: pH 7.9, 10 mM KCl, 2 mM MgCl_2_, 0.1 mM EDTA, 0.2% NP40) with protease inhibitor (#11697498001, Roche). The supernatant (cytoplasmic fraction) was collected after centrifuged at 13,000 rpm at 4°C for 10 min. After washed twice, the nuclei pellet was further lysed with 250 μl × #9803 cell lysis buffer with protease inhibitor. The nuclear fraction was collected after centrifuged at 13,000 rpm at 4°C for 10 min after sonication. The subfraction samples were subjected to immunoblotting analysis.

### Luciferase assay

Cells were transfected with indicated plasmid using electroporation, and pRL-TK was used as an internal control. After cultured for 24 hours, cells were treated with trametinib for another 36 hours. After treatment stopped, relative luciferase units (RLU) were measured using the Dual-Glo Luciferase Assay System (Promega) according to the manufacturer’s instructions. RLUs from firefly luciferase signal were normalized by RLUs from Renilla signal.

### Flow cytometric analysis

After treated with 100 nM trametinib or 0.3% DMSO for 48 hours, cells were fixed with 1 mL of 70% ice-cold ethanol at 4°C overnight. After washed with 1× PBS, cells were stained with propidium iodide (C1052, Beyotime) at 37°C for 30 min and were analyzed using Beckman Gallios flow cytometry. Data were analyzed using a FlowJo software.

### GeCKO library screen

The complete GeCKO library screen procedure was previously described (Liu et al, 2019). Briefly, GSU cells were infected with an optimal volume of viruses containing sgRNA library at an MOI of 0.4, and were subjected to puromycin for 7 days to remove uninfected cells. After puromycin selection, 2×10 ^7^ GSU cells were collected as DMSO_Day0_ group and were stored at −80°C. The remaining cells were divided into two groups, followed by 0.015% DMSO (DMSO_Day14_ group) or 15 nM trametinib for another 14 days (Trametinib_Day14_ group). To keep 300-fold sgRNA coverage, 2-3 ×10^7^ living cells were harvested for each group when treatment stopped. The genomic DNA was extracted with Qiagen Blood & Cell Culture Midi kit, followed by PCR amplification, and was sequenced using Illumina HiSeq 2500 by GENEWIZ (Suzhou, China).

### *In vivo* tumour xenograft models

BALB/c nude mice were purchased from Shanghai SLAC Laboratory Animal Company. All the experimental procedures were approved by the Institutional Animal Care and Use Committee of Shanghai Institute for Nutrition and Health. Cells were injected subcutaneously in the right flank of each mouse. Mice were divided into different groups based on the average tumour volume of each group, approximately 100-150 mm^3^. Trametinib (3 mg/kg), or vehicle (10% PEG400, 10% Cremophor® EL) was orally administrated, twice per week. Doxycycline (1 mg/ml) was added to the drinking water to induce p53 expression. Then tumour was measured twice weekly and tumour volume was calculated by the formula V= 0.5×L×W ^2^ (V: volume, L: length, W: width).

### Statistical analysis

Data represent means ±SD of 2-3 independent repeats. Statistical significance was determined by Student’s *t*-test and different levels of statistical significance were denoted by p-values (*p < 0 .05, **p < 0 .01, ***p <0.001).

## Supporting information

F:\&#30740;&#31350;&#29983;\p53\puma\xw and xq\submit 20210714

## Acknowledgments

This work is financially supported by grants from Chinese Academy of Sciences (QYZDB-SSW-SMC034, XDA12020210), National Natural Science Foundation of China (U210220136, 81322034) and National Key R&D Program of China (2016YFC1302400).

We thank Ms. Xu Jiang and Mr. Zizhuo Chen for constructing *TP53* mutant plasmids, and thank the excellent service of institutional core facilities including Molecular Biology and Biochemistry and laboratory animal technical platforms.

## Declarations of interests

The authors declare that they have no conflicts of interest.

## Author Contributions

J.Y.L. conceived and supervised the project. J.Y.L., X.W. and Q.X. wrote the manuscript and the other authors revised it. X.W. and X.Q. performed most of the experiments. J.Y.S. did immunofluorescent and biochemical experiments. Y.J. performed the bioinformatica analyses. Y.F.Z. performed cell viability assay and cell culture. X.Y.K., F.J., L.Z. and C.Q. helped the improvement of the project and extensive discussions.

## Availability of data

The GeCKO library data has been deposited in the NCBI GEO (http://www.ncbi.nlm.nih.gov/geo) under accession number GSE152846. All relevant data supporting the key findings are available from the corresponding author upon reasonable request.

## Notes

### Competing Interest Statement

The authors have declared no competing interest.

