## Supplementary material for "Targeting KRAS-mutant stomach/colorectal tumours by disrupting the ERK2-p53 complex": F:\&#30740;&#31350;&#29983;\p53\puma\xw and xq\submit 20210714

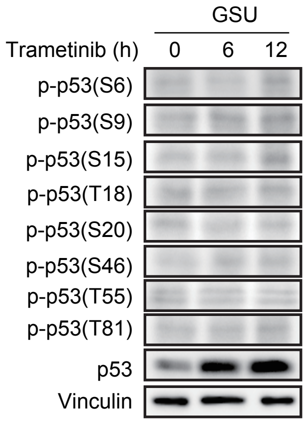

**Supplemental Figure 1.** p53 phosphorylation is not detectable in the transactivation domain before and after trametinib treatment. The phosphorylation sites of the p53 TAD were determined after treatment with 100 nM trametinib for the indicated times.

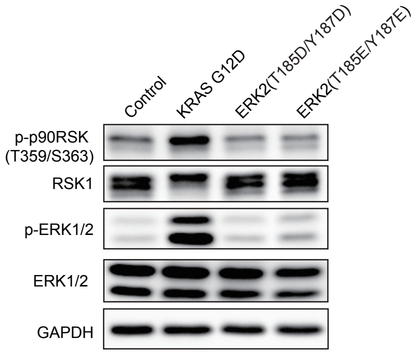

**Supplemental Figure 2.** p53 T185D/Y187D and T185E/Y187E mutants failed to induce ERK substrate p90RSK phosphorylation. Indicated protein expression levels were determined by immunoblottoing after vector control, KRAS G12D, ERK2 T185D/Y187D and T185E/Y187E mutant were introduced into 293T cells. GAPDH was used as a loading control.

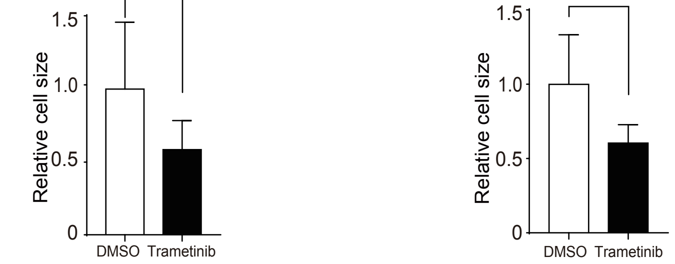

**Supplemental Figure 3.** Trametinib significantly reduced the cell size of AGS and GSU cells by approximately 50%. The relative length and width of AGS and GSU cells were determined by scale bar under confocal microscopy after trametinib treatment for 24 hous. Cell area = length x width. At least 50 cells were counted in each group. Data represent mean ± SD of 2-3 independent expriments. ****p<0.0001, by Student's t-test.

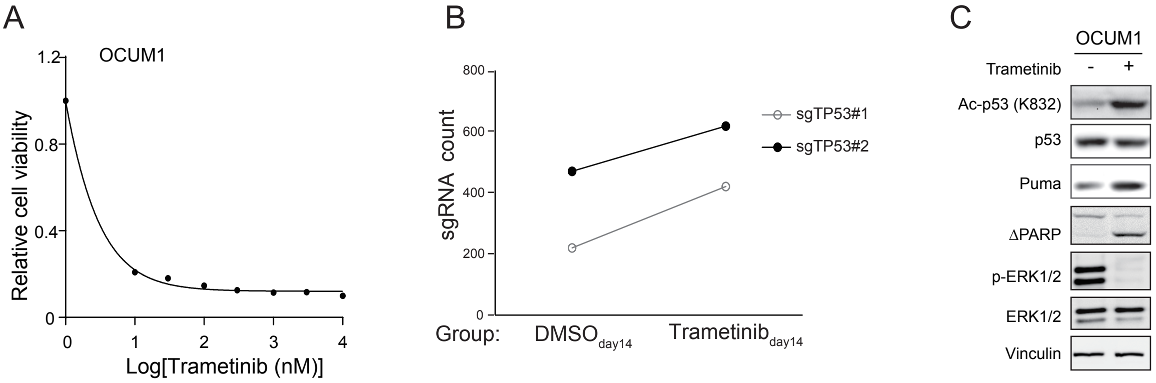

**Supplemental Figure 4.** Genome-wide knockout screening identifies that *TP53* is critical for trametinib-induced apoptosis of MAP2K1$^{Q56P}$ mutant OCUM1 cells. (A) MAP2K1$^{Q56P}$ mutant OCUM1 were sensitive to trametinib with an $IC_{50}$ value of 5 nM. The cell viability was determined at $OD_{570}$ by an MTT assay; the data represent the means ± SD of 3 independent experiments. (B) Genome-scale screening revealed that the levels of sgRNAs targeting the *TP53* gene were increased approximately1.3- and 1.9-fold in the Trametinib$_{day14}$ group compared to the DMSO$_{day14}$ group (FDR<0.05). (C) The expression levels of the indicated proteins were detected in the absence or presence of 100 nM trametinib by immunoblotting. Vinculin was used as a loading control.

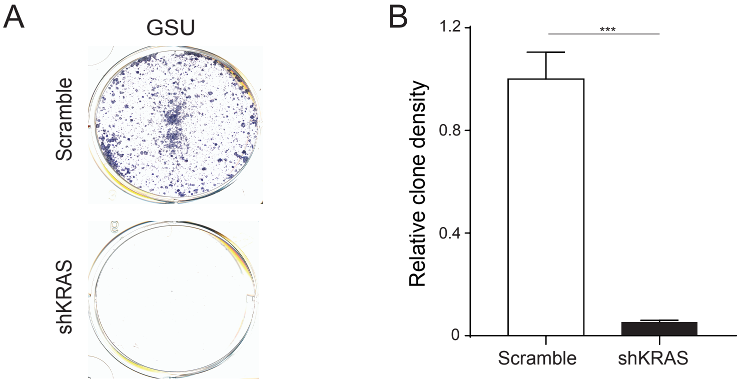

**Supplemental Figure 5.** KRAS is critical for maintaining the survival of KRAS$^{G12D}$ mutant GSU cells. (A) 1,000 GSU cells with scramble shRNA or shKRAS knockdown were seeded into each well of a 6-well plate, and stained with 0.5% crystal violet after two weeks of culture. (B)The colony density of each well was captured and calculated with ImageJ software. Data represent mean ± SD of 3 independent expriments. ***p<0.001, by Student's t-test.

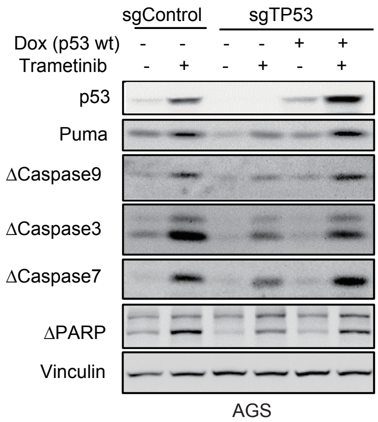

**Supplemental Figure 6.** Trametinib promotes the apoptosis of KRAS$^{G12D}$ mutant AGS cells in a p53-dependent manner. Indicated proteins were determined by immunoblotting after treated with 100 nM trametinib for 24 hours in sgControl, sgTP53 and sgTP53 with reintroduced wild-type p53 AGS cells.

A

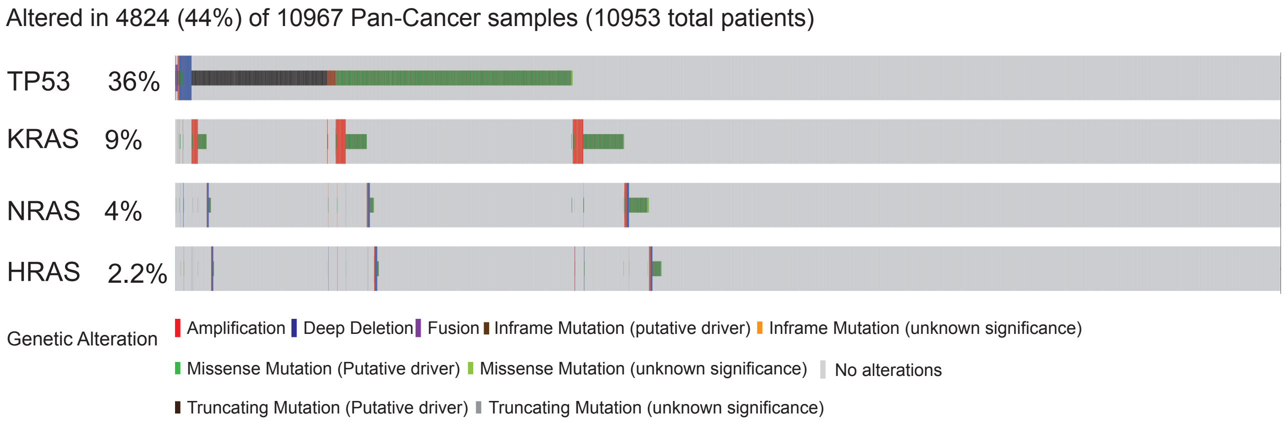

B

| TCGA PanCancer Atlas (Stomach) | | | |
|---|---|---|---|
| p=0.3648 | | KRAS | |
| | | Mutant | Wild-type |
| TP53 | Mutant | 31 | 183 |
| | Wild-type | 39 | 181 |

| Stomach Cancer (Onco SG 2018) | | | |
|---|---|---|---|
| p>0.9999 | | *KRAS* | |
| | | Mutant | Wild-type |
| TP53 | Mutant | 8 | 44 |
| | Wild-type | 9 | 47 |

C

| TCGA PanCancer Atlas (Colorectal) | | | |
|---|---|---|---|
| p=0.5268 | | KRAS | |
| | | Mutant | Wild-type |
| TP53 | Mutant | 127 | 186 |
| | Wild-type | 93 | 120 |

| Colorectal cancer(DFCI, Cell reports, 2016) | | | |
|---|---|---|---|
| p=0.2096 | | KRAS | |
| | | Mutant | Wild-type |
| TP53 | Mutant | 82 | 238 |
| | Wild-type | 91 | 208 |

D

| Colorectal cancer (MSKCC, Cancer Cell, 2018) | | | |
|---|---|---|---|
| p<0.0001 | | KRAS | |
| | | Mutant | Wild-type |
| TP53 | Mutant | 100 | 527 |
| | Wild-type | 200 | 307 |

**Supplemental Figure 7.** *KRAS* is frequently co-mutated with *TP53* in the Pan-cancer Atlas samples. (A) TCGA mutation data analysis revealed that TP53 mutations occurred in 479 out of 971 KRAS-mutant tumour samples (co-occurrence, p<0.001), 140 out of 376 NRAS-mutant tumour samples (mutual exclusivity, p=0.539), and 91 out of 227 HRAS-mutant tumour samples (co-occurrence, p=0.539). (B) TP53 mutations was not significantly correlated with KRAS mutations in stomach cancer samples. (C) TP53 mutations was not significantly correlated with KRAS mutations in colorectal cancer samples. (D) TP53 wild-type were predominant in KRAS-mutant metastatic colorectal cancer samples compared to that of TP53 mutatations. Mutation data were obtained from the TCGA database via the cBioPortal website. The statistical analysis was performed with two sided Fisher's exact test.

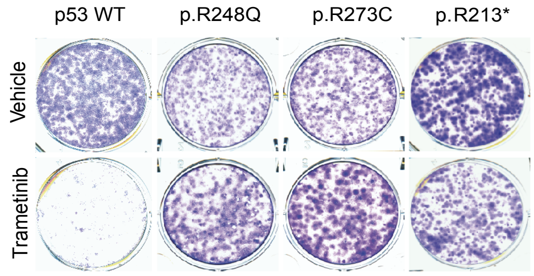

**Supplemental Figure 8.** TP53 mutations occurred in the DNA binding domain (amino acids 245-285) and nonsense mutations confered resistance to trametinib in KRAS-mutant tumour cells. TP53 knockout AGS cells were reintroduced with the indicated TP53 mutants and subjected to a clonogenicity assay after treatment with 1 nM trametinib for 14 days. Wild-type p53 was used as a positive control.

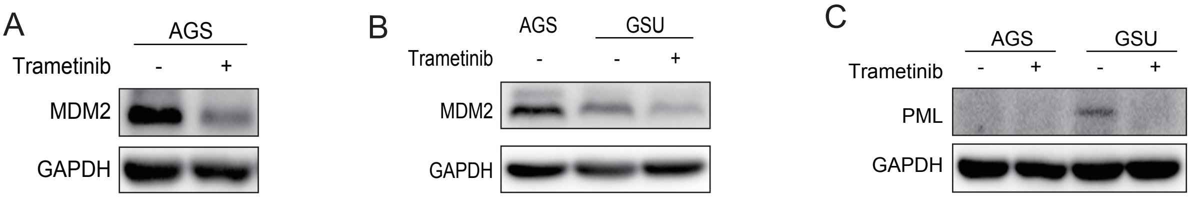

**Supplemental Figure 9.** Trametinib reduced MDM2 expression in KRAS-mutant stomach/colorectal tumor cells. (A) MDM2 protein expression was determined in AGS cells by immunoblotting after treatment with 1 μM trametinib for 24 hours. GAPDH was used as a loading control. (B) MDM2 protein expression was determined by immunoblotting after treatment with 1 μM trametinib for 12 hours. AGS cells were used as a positive control. (C) PML has little or no expression in KRAS-mutant GSU and AGS cell lines, which was further inhibited by trametinib treatment.

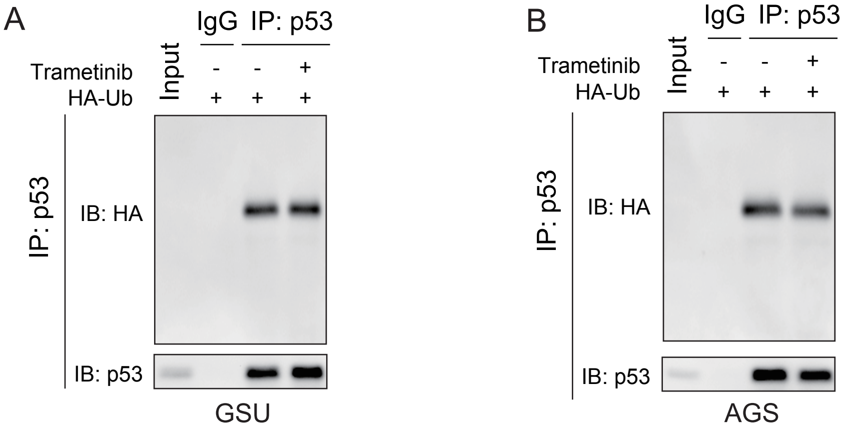

**Supplemental Figure 10.** Trametinib didn't affect the ubiquitination of p53 protein. (A) Trametinib had no effect on the ubiquitination of p53 protein in GSU cells. The protein expression levels were determined by immunoblotting after immunopreciptated with p53 antibody when GSU cells were transfected with HA-Ub for 48 hours. (B) Trametinib had no effect on the ubiquitination of p53 protein in AGS cells. The protein expression levels were determined by immunoblotting after immunopreciptated with p53 antibody when AGS cells were transfected with HA-Ub for 48 hours.

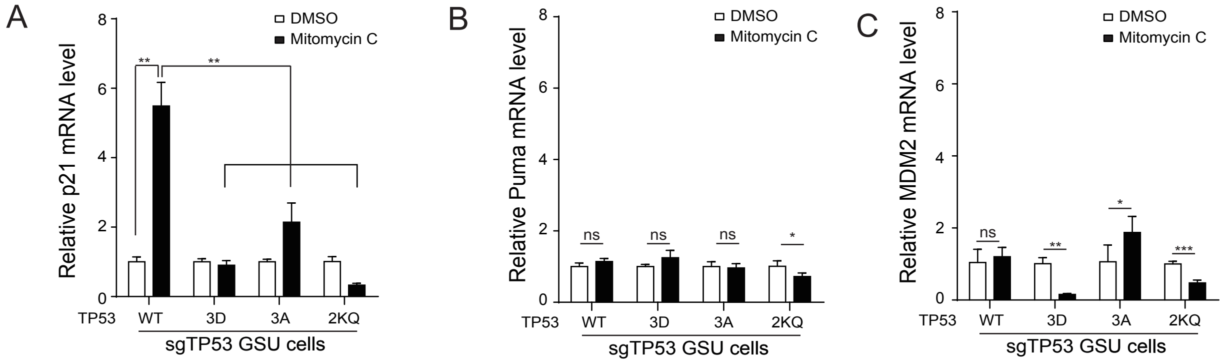

**Supplemental Figure 11.** Acetylated p53 is not responding to mitomycin C for inducing p21 expression. (A-C) The p21, Puma and MDM2 mRNA expression levels were determined by qRT-PCR when indicated p53 wild-type and mutants were re-introduced back into sgTP53 GSU cells following treatment of 1 μM mitomycin C for 36 hours.

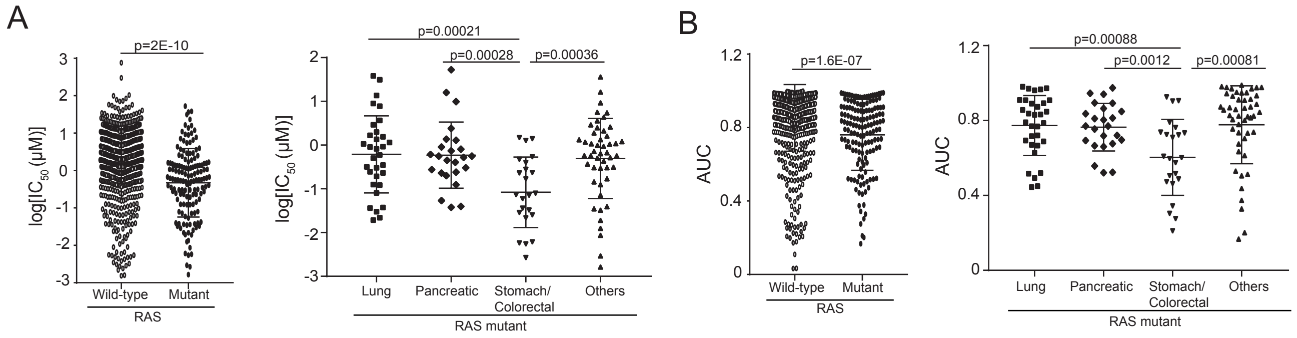

**Supplemental Figure 12.** Trametinib preferentially induces the apoptosis of KRAS-mutant stomach/colorectal cancer cell lines. (A-B) The half maximal inhibitory concentrations ($IC_{50}$) and area under the curve (AUC) of 771 human cancer cell lines responding to trametinib were obtained from the Genomics of Drug Sensitivity in Cancer database. The statistical significance was calculated by Student's t-test after they were divided into two groups (RAS wild-type, n=623; RAS mutant, n=148). Cancer cell lines harbouring RAS hotspot mutations at glycine 12, glycine 13 and glutamine 61 sites were further divided into four subgroups, including RAS-mutant lung cancer (n=32), RAS-mutant pancreatic cancer (n=24), RAS-mutant stomach/colorectal cancer (n=23) and other RAS-mutant cancers (n=48). Twenty-one cell lines with non-hotspot RAS mutations were excluded.

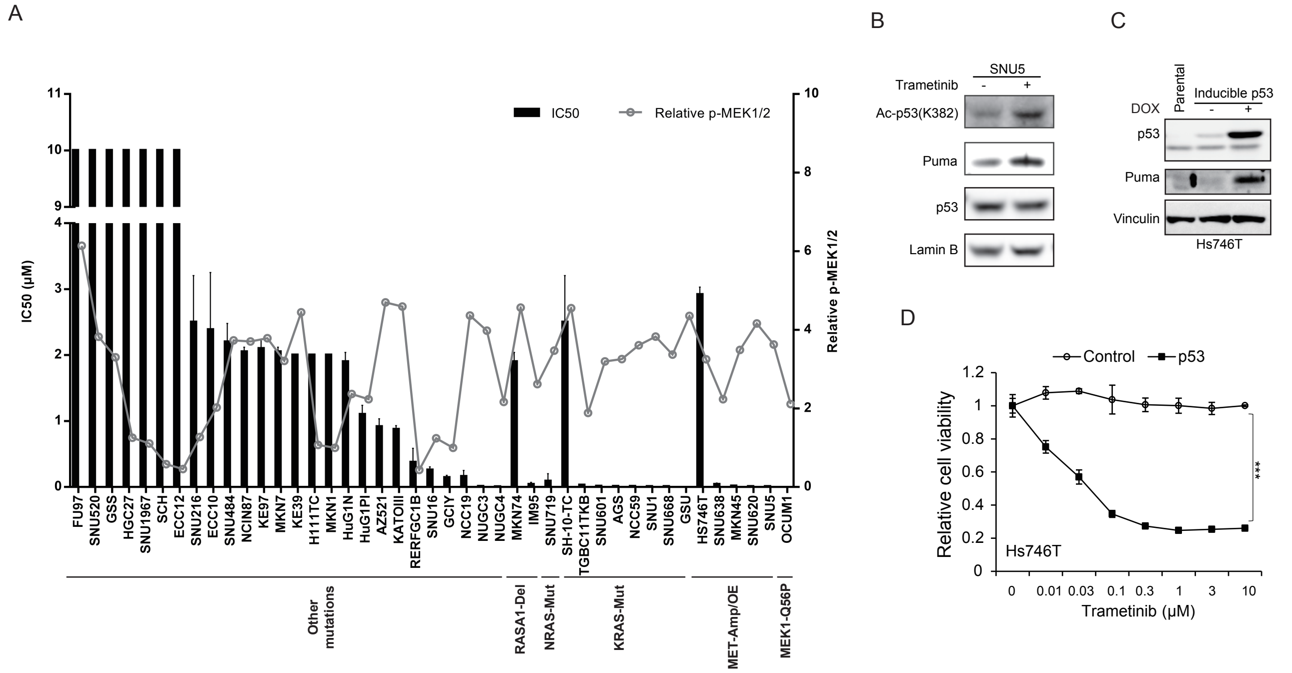

**Supplemental Figure 13.** Stomach cancer cell lines harbouring KRAS, MAP2K1 and MET mutations were sensitive to trametinib treatment. (A) Trametinib potently inhibited the viability of 18 of 33 MEK1/2 -activated stomach cancer cell lines harboring KRAS, NRAS, RASA1, MET and MAP2K1 mutations. The levels of phosphorylated MEK1/2 was detected under serum-free condition using immunoblotting and was calculated and presented on the right. (B) MET-amplified Hs746T cells were primary resistant to trametinib due to the $TP53^{K319*}$ nonsense mutation, and were resensitiezed to trametinib when wild-type p53 was put back. The viability was determined by an MTT assay. (C) The protein expression levels of p53 and Puma in Hs746T cells with or without wild-type p53 expression were detected by immunoblotting. (D) The protein expression levels of indicated proteins in MET-amplified SNU5 cells were detected by immunoblotting after 100 nM trametinib treatment. Data represent means ± SD of 3 independent experiments. ***p<0.001, by Student's t-test.

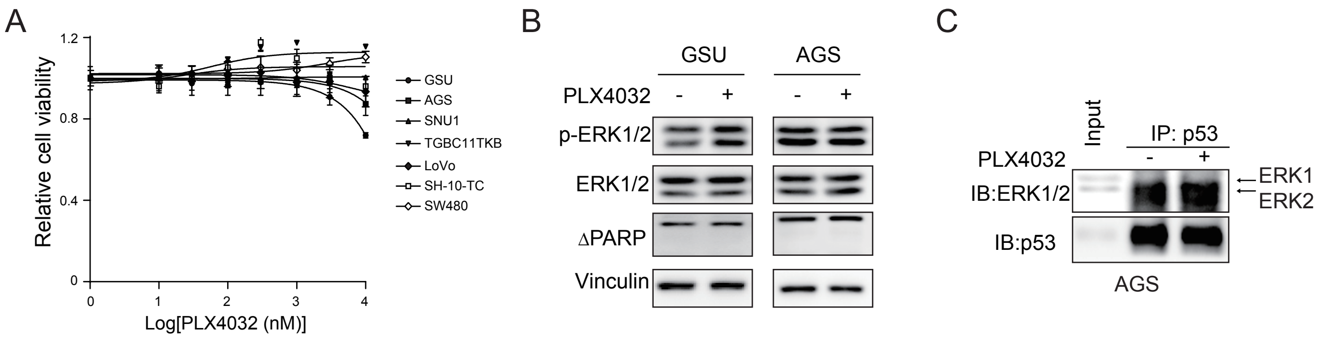

**Supplemental Figure 14.** PLX4032 has no inhibitory effects on KRAS-mutant tumour cells due to its inability to disrupt the ERK2-p53 complex. (A) The viability of 7 KRAS-mutant stomach/colorectal cancer cell lines treated with PLX4032 for 72 hours at the indicated concentrations were determined. (B) PLX4032 did not reduce ERK1/2 phosphorylation in either $KRAS^{G12D}$ mutant GSU or AGS cell lines. The protein expression levels were determined by immunoblotting after treatment with 100 nM PLX4032 for 12 hours. (C) PLX4032 did not affect the binding of ERK2 to p53 protein. Co-IP was used to detect the binding affinity between ERK2 and p53 in the absence or presence of PLX4032.

**Table S1. The top 25 ranked genes of which sgRNAs were enriched in $\text{Trametinib}_{day14}/\text{DMSO}_{day14}$ group (A)**

| Gene | crisprGeneScore | Rank |
|---|---|---|
| ALB | 2.661719 | 1 |
| MYH9 | 2.64502 | 2 |
| RPS10-NUDT3 | 2.503342 | 3 |
| KRAS | 2.48668 | 4 |
| POLR3H | 2.377377 | 5 |
| AP2S1 | 2.180823 | 6 |
| MYL6 | 2.125992 | 7 |
| RPL14 | 2.075375 | 8 |
| ORM1 | 2.031758 | 9 |
| HP | 1.998332 | 10 |
| MED23 | 1.990867 | 11 |
| RPL3 | 1.987171 | 12 |
| YARS | 1.937788 | 13 |
| CCNA2 | 1.908202 | 14 |
| RPL11 | 1.846019 | 15 |
| ROCK2 | 1.795084 | 16 |
| FAM96B | 1.773304 | 17 |
| POLR3K | 1.758479 | 18 |
| BCL9L | 1.745305 | 19 |
| PRPF38B | 1.745224 | 20 |
| RIMBP3B | 1.739872 | 21 |
| **TP53** | **1.72739** | **22** |
| RPS19 | 1.716801 | 23 |
| ARL17A | 1.709374 | 24 |
| EIF1AX | 1.690801 | 25 |

**Table S1. The top 25 ranked genes of which sgRNAs were decreased in Trametinib$_{day14}$/DMSO$_{day14}$ group (B)**

| Gene | crisprGeneScore | Rank |
|---|---|---|
| HNRNPLL | -1.29226 | 1 |
| EPPIN-WFDC6 | -1.18432 | 2 |
| RLN2 | -1.16077 | 3 |
| SRPX | -1.1245 | 4 |
| ZADH2 | -1.11766 | 5 |
| PURG | -1.10906 | 6 |
| NBPF4 | -1.0706 | 7 |
| HIST2H2AA4 | -1.0643 | 8 |
| RBMY1F | -1.05752 | 9 |
| OR4A5 | -1.05437 | 10 |
| FAM47A | -1.0269 | 11 |
| ZGPAT | -1.02419 | 12 |
| GLCE | -0.97781 | 13 |
| JSRP1 | -0.97505 | 14 |
| HYKK | -0.97052 | 15 |
| HNRNPH2 | -0.96216 | 16 |
| PTPLA | -0.95792 | 17 |
| C8orf42 | -0.93915 | 18 |
| APOBEC4 | -0.93445 | 19 |
| STON1-GTF2A1L | -0.91396 | 20 |
| VPS8 | -0.9137 | 21 |
| STAG2 | -0.90497 | 22 |
| AAED1 | -0.89844 | 23 |
| MPZL1 | -0.89491 | 24 |
| PIRT | -0.89224 | 25 |

**Table S1. The top 25 ranked genes of which sgRNAs were enriched in $DMSO_{day14}$/$DMSO_{day0}$ group (C)**

| Gene | crisprGeneScore | Rank |
|---|---|---|
| AJUBA | 1.896865 | 1 |
| NBPF6 | 1.309073 | 2 |
| PAK3 | 0.989973 | 3 |
| PTPN14 | 0.9501 | 4 |
| POTEH | 0.94905 | 5 |
| KSR2 | 0.936368 | 6 |
| APOBEC4 | 0.934177 | 7 |
| C7orf72 | 0.929174 | 8 |
| OCA2 | 0.922727 | 9 |
| OR4A5 | 0.891062 | 10 |
| MBD3L5 | 0.877181 | 11 |
| PTPLA | 0.861795 | 12 |
| PACSIN2 | 0.856736 | 13 |
| VCY1B | 0.854591 | 14 |
| RAD17 | 0.851966 | 15 |
| KIAA0232 | 0.849355 | 16 |
| RBMY1F | 0.839436 | 17 |
| KDM5D | 0.839268 | 18 |
| RHBDF2 | 0.836334 | 19 |
| RGS11 | 0.828824 | 20 |
| SERGEF | 0.826061 | 21 |
| PIRT | 0.825974 | 22 |
| PSG4 | 0.817532 | 23 |
| PCDHGC5 | 0.810051 | 24 |
| HNRNPLL | 0.809612 | 25 |

**Table S1. The top 25 ranked genes of which sgRNAs were decreased in $DMSO_{day14}/DMSO_{day0}$ group (D)**

| Gene | crisprGeneScore | Rank |
|---|---|---|
| FAM86B1 | -2.55202 | 1 |
| RPS10-NUDT3 | -2.32642 | 2 |
| YARS | -2.17135 | 3 |
| RPL14 | -2.11751 | 4 |
| RPL3 | -2.11233 | 5 |
| SBNO1 | -2.10867 | 6 |
| POLR3H | -2.04766 | 7 |
| MKI67IP | -1.9707 | 8 |
| SPC24 | -1.93943 | 9 |
| POP5 | -1.93431 | 10 |
| ARL17A | -1.90627 | 11 |
| WASH1 | -1.88631 | 12 |
| PHF5A | -1.87817 | 13 |
| RPL34 | -1.86797 | 14 |
| BCL9L | -1.84293 | 15 |
| NIP7 | -1.80432 | 16 |
| RPL11 | -1.79601 | 17 |
| RPS19 | -1.78865 | 18 |
| POLR3K | -1.75791 | 19 |
| MMS22L | -1.73542 | 20 |
| TRIP13 | -1.72797 | 21 |
| VARS | -1.70415 | 22 |
| OGFOD2 | -1.65493 | 23 |
| RBM14 | -1.65476 | 24 |
| MRPL33 | -1.65005 | 25 |

**Table S2. Primers for constructing TP53 and ERK2 mutants**

| Primer Name | Sequence（5' to 3'） |
|---|---|
| Forward primer for TP53 cDNA | CGGAATTCCGatggaggagccgca |
| Reverse primer for TP53 cDNA | AAATATGCGGCCGCtcagtctgagt |
| Forward primer for pENTR-HA-p53 | GATCCatgtacccctacgacgtccccgactacgccG |
| Reverse primer for pENTR-HA-p53 | AATTCggcgtagtcggggacgtcgtaggggtacatG |
| Forward primer 1 for p53 mutant plasmids | CGCGGATCCATGTACCCCTACGACGTC |
| Reverse primer 1 for p53 mutant plasmids | CCGCTCGAGTTACTATCAGTCTGAGTCAG |
| Forward primer 2 for p53 V157F mutant plasmid | ACGTCTCActtcCGCGCCATGGCCATCTACAAGCAGTCACAGCACATGA |
| Reverse primer 2 for p53 V157F mutant plasmid | ACGTCTCAgaagCGGGTGCCGGGCGGGGGTGTGGAATCAACCCACAGCTGCAC |
| Forward primer 2 for p53 Y163C mutant plasmid | ACGTCTCAgcaaGCAGTCACAGCACATGACGGAGGTTGTGA |
| Reverse primer 2 for p53 Y163C mutant plasmid | ACGTCTCAttgcAGATGGCCATGGCGCGGACGCGGGTG |
| Forward primer 2 for p53 R175H mutant plasmid | ACGTCTCAgaggCACTGCCCCCACCATGAGCGCTGCTCAGATAG |
| Reverse primer 2 for p53 R175H mutant plasmid | ACGTCTCAcctcACAACCTCCGTCATGTGCTGTGACTGCTTG |
| Forward primer 2 for p53 C176F mutant plasmid | ACGTCTCAgaggCGCTtCCCCCACCATGAGCGCTGCT |
| Reverse primer 2 for p53 C176F mutant plasmid | ACGTCTCAcctcACAACCTCCGTCATGTGCTGTGACTGCTTG |
| Forward primer 2 for p53 H179R mutant plasmid | ACGTCTCAccacCGTGAGCGCTGCTCAGATAGCGAT |

| | |
|---|---|
| Reverse primer 2 for p53 H179R mutant plasmid | ACGTCTCAgtggGGGCAGCGCCTCACAACCTCCGT |
| Forward primer 2 for p53 H179Y mutant plasmid | ACGTCTCAccactATGAGCGCTGCTCAGATAGCGAT |
| Reverse primer 2 for p53 H179Y mutant plasmid | ACGTCTCAgtggGGGCAGCGCCTCACAACCTCCGT |
| Forward primer 2 for p53 H193R mutant plasmid | ACGTCTCAcagcgTCTTATCCGAGTGGAAGGAAATTT |
| Reverse primer 2 for p53 H193R mutant plasmid | ACGTCTCAgctgAGGAGGGGCCAGACCATCGCTATCTGA |
| Forward primer 2 for p53 I195T mutant plasmid | ACGTCTCAcagcATCTTAcCCGAGTGGAAGGAAATTTGCGTGT |
| Reverse primer 2 for p53 I195T mutant plasmid | ACGTCTCAgctgAGGAGGGGCCAGACCATCGCTATCTGA |
| Forward primer 2 for p53 R213* mutant plasmid | ACGTCTCAttttGACATAGTGTGGTGGTGCCCTATGA |
| Reverse primer 2 for p53 R213* mutant plasmid | ACGTCTCAaaaaGTGTTTCTGTCATCCAAATACTCCACACGCAAATTTCCTTCCAC |
| Forward primer 2 for p53 Y220C mutant plasmid | ACGTCTCAttttCGACATAGTGTGGTGGTGCCCTGTGAGCCGCCTGAGGTT |
| Reverse primer 2 for p53 Y220C mutant plasmid | ACGTCTCAaaaaGTGTTTCTGTCATCCAAATACTCCACACGCAAATTTCCTTCCAC |
| Forward primer 2 for p53 Y234C mutant plasmid | ACGTCTCAcactgCAACTACATGTGTAACAGTTCCTGCAT |
| Reverse primer 2 for p53 Y234C mutant plasmid | ACGTCTCAagtgGATGGTGGTACAGTCAGAGCCAACCTCA |
| Forward primer 2 for p53 S241F mutant plasmid | ACGTCTCAcactACAACTACATGTGTAACAGTTtCTGCATGGGCGGCATGAA |
| Reverse primer 2 | ACGTCTCAagtgGATGGTGGTACAGTCAGAGCCAACCTCA |

| | |
|---|---|
| for p53 S241F mutant plasmid | |
| Forward primer 2 for p53 G245S mutant plasmid | ACGTCTCAgggcaGCATGAACCGGAGGCCCATCCTCA |
| Reverse primer 2 for p53 G245S mutant plasmid | ACGTCTCAgcccATGCAGGAACTGTTACACATGTAGTT |
| Forward primer 2 for p53 G245D mutant plasmid | ACGTCTCAgggcGaCATGAACCGGAGGCCCATCCTCA |
| Forward primer 2 for p53 R248Q mutant plasmid | ACGTCTCAgggcGGCATGAACCaGAGGCCCATCCTCACCATCATCA |
| Forward primer 2 for p53 R248W mutant plasmid | ACGTCTCAgggcGGCATGAACtGGAGGCCCATCCTCACCATCATCA |
| Forward primer 2 for p53 R248L mutant plasmid | ACGTCTCAgggcGGCATGAACCtGAGGCCCATCCTCACCATCATCA |
| Forward primer 2 for p53 R249S mutant plasmid | ACGTCTCAgggcGGCATGAACCGGAGtCCCATCCTCACCATCATCA |
| Forward primer 2 for p53 V272M mutant plasmid | ACGTCTCAtgagaTGCGTGTTTGTGCCTGTCCTGGGAGAGA |
| Reverse primer 2 for p53 V272M/R273H/R273C/R273L mutant plasmids | ACGTCTCActcaAAGCTGTTCCGTCCCAGTAGA |
| Forward primer 2 for p53 R273H mutant plasmid | ACGTCTCAtgagGTGCaTGTTTGTGCCTGTCCTGGGAGAGA |
| Forward primer 2 for p53 R273C mutant plasmid | ACGTCTCAtgagGTGtGTGTTTGTGCCTGTCCTGGGAGAGA |
| Forward primer 2 for p53 R273L mutant plasmid | ACGTCTCAtgagGTGCtTGTTTGTGCCTGTCCTGGGAGAGA |
| Forward primer 2 for p53 R282W mutant plasmid | ACGTCTCAagactGGCGCACAGAGGAAGAGAATCTCCGCAA |
| Reverse primer 2 | ACGTCTCAgtctCTCCCAGGACAGGCACAAA |

| | |
|---|---|
| for p53 R282W mutant plasmid | |
| Forward primer 2 for p53 E285K mutant plasmid | ACGTCTCAagacCGGCGCACAaAGGAAGAGAATCTCCGCAA |
| Reverse primer 2 for p53 E285K mutant plasmid | ACGTCTCAgtctCTCCCAGGACAGGCACAAA |
| Forward primer 2 for p53 5KQ/5KR/2KQ/2KR/K382Q/K382R mutant plasmids | CTTTGAGGTGCGTGTTTGTG |
| Reverse primer 2 for p53 5KQ mutant plasmid | TAGATATCAGTCTGAGTCAGGCCCTTCTGTCTTGAACATGAGTTgTTgATGGCGactaGTAGACTGACCCTgTTgGGACTgCAGGTGGCTGGAGT |
| Reverse primer 2 for p53 5KR mutant plasmid | TAGATATCAGTCTGAGTCAGGCCCTTCTGTCTTGAACATGAGTcTTcTATGGCGactaGTAGACTGACCCcTTcTGGACcTCAGGTGGCTGGAGT |
| Reverse primer 2 for p53 2KQ/2KR mutant plasmids | TGGCGactaGTAGACTGACCCTTTTTGGACTTCAGGTG |
| Reverse primer 2 for p53 K382Q mutant plasmid | TAGATATCAGTCTGAGTCAGGCCCTTCTGTCTTGAACATGAGTTgTTTATGGCGactaGTAGACTGACCCTTTTTGGACTTCAGGTG |
| Reverse primer 2 for p53 K382R mutant plasmid | TAGATATCAGTCTGAGTCAGGCCCTTCTGTCTTGAACATGAGTcTTTTATGGCGactaGTAGACTGACCCTTTTTGGACTTCAGGTG |
| Forward primer 2 for p53 K120Q/K120R mutant plasmids | aCGTCTCgtctgtgacttgcacgtactcccctgccctcaac |
| Reverse primer 2 for p53 K120Q mutant plasmid | aCGTCTCacagactgggctgtcccagaatgcaagaagcccagac |
| Reverse primer 2 for p53 K120R mutant plasmid | aCGTCTCacagacctggctgtcccagaatgcaagaagcccagac |
| Forward primer 2 for KRAS G12D mutant plasmid | GAGCTCGGATCCATGACTGAATATAAACTTGTGG |
| Reverse primer 2 for KRAS G12D | GCATGCTCGAGTTACATTATAATGCATTTTTTAATTTTC |

| | |
|---|---|
| mutant plasmid | |
| Forward primer 2 for MEK1 Q56P mutant plasmid | aaaGGATCCcccaagaagaagccgac |
| Reverse primer 2 for MEK1 Q56P mutant plasmid | agatctagacgccagcagcatgggttggtgtgctgggctggttaaggccgatggtggag |
| Forward primer for ERK2 T185A Y187F/T185D Y187D/T185E Y187E/T185A Y187D/T185D Y187F mutant plasmids | CGATTACAAGGATGACGATGACAAGC |
| Reverse primer 1 for ERK2 T185A Y187F mutant plasmid | cctgtaccaacgtgtggccacaaattccgccaggaaccctgtgtgatcatg |
| Reverse primer 1 for ERK2 T185D Y187D mutant plasmid | cctgtaccaacgtgtggccacatcttcatccaggaaccctgtgtgatcatg |
| Reverse primer 1 for ERK2 T185E Y187E mutant plasmid | cctgtaccaacgtgtggccacttcttcttccaggaaccctgtgtgatcatg |
| Reverse primer 1 for ERK2 T185A Y187D mutant plasmid | cctgtaccaacgtgtggccacatcttccgccaggaaccctgtgtgatcatg |
| Reverse primer 1 for ERK2 T185D Y187F mutant plasmid | cctgtaccaacgtgtggccacaaattcatccaggaaccctgtgtgatcatg |
| Reversr primer 2 for ERK2 T185A Y187F/T185D Y187D/T185E Y187E/T185A Y187D/T185D Y187F mutant plasmids | ccggaattcaacataatttctggagccctgtaccaacgtgtggccac |

**Table S3. Oligos for sgRNAs and shRNAs**

| primer name | Sequence（5' to 3'） |
|---|---|
| shKRAS-G12D | CCGGGTTGGAGCTGATGGCGTAGCTCGAGCTACGCCATCAGCTCCAACTTTTTG |
| shERK1#1 | CCGGGCCATGAGAGATGTCTACACTCGAGTGTAGACATCTCTCATGGCTTTTTG |
| shERK1#2 | CCGGTCCCTGTCAAAGCTGTCACTTCTCGAGAAGTGACAGCTTTGACAGGGATTTTTG |
| shERK2#1 | CCGGGAGGATTGAAGTAGAACAGCTCGAGCTGTTCTACTTCAATCCTCTTTTTG |
| shERK2#2 | CCGGCAAAGTTCGAGTAGCTATCAACTCGAGTTGATAGCTACTCGAACTTTGTTTTTG |
| shPUMA #1 | CCGGCGTGAAGAGCAAATGAGCCAACTCGAGTTGGCTCATTTGCTCTTCACGTTTTTG |
| shPUMA #2 | CCGGGAGGGTCCTGTACAATCTCATCTCGAGATGAGATTGTACAGGACCCTCTTTTTG |
| shPUMA #3 | CCGGACGGTCCTCAGCCCTCGCTCTCTCGAGAGAGCGAGGGCTGAGGACCGTTTTTTG |
| sgTP53-1 | CACCGCCGGTTCATGCCGCCCATGC |
| sgTP53-2 | CACCGCGCTATCTGAGCAGCGCTCA |
| sgTP53-3 | CACCGCCCCTTGCCGTCCCAAGCAA |
| sgEGFP | CACCGGAGCTGGACGGCGACGTAAA |

**Table S4. Primers for Real time PCR**

| gene name | forward primer | reverse primer |
|---|---|---|
| TP53 | GCCCAACAACACCAGCTCCT | CCTGGGCATCCTTGAGTTCC |
| GAPDH | GAGTCAACGGATTTGGTCGT | GACAAGCTTCCCGTTCTCAG |
| MDM2 | GGACTCGGAAGATTACAGCCTGA | TGTCTGATAGACTGTGACCCG |
| P21 | AGATCCACAGCGATATCCAGAC | ACCGAAGAGACAACGGCACACT |
| PUMA | TTGTGCTGGTGCCCGTTCCA | AGGCTAGTGGTCACGTTTGGCT |
| BAX | CAGGATGCGTCCACCAAGAA | AGTCCGTGTCCACGTCAGCA |
